# Structures of the Omicron spike trimer with ACE2 and an anti-Omicron antibody: mechanisms for the high infectivity, immune evasion and antibody drug discovery

**DOI:** 10.1101/2021.12.27.474273

**Authors:** Wanchao Yin, Youwei Xu, Peiyu Xu, Xiaodan Cao, Canrong Wu, Chunyin Gu, Xinheng He, Xiaoxi Wang, Sijie Huang, Qingning Yuan, Kai Wu, Wen Hu, Zifu Huang, Jia Liu, Zongda Wang, Fangfang Jia, Kaiwen Xia, Peipei Liu, Xueping Wang, Bin Song, Jie Zheng, Hualiang Jiang, Xi Cheng, Yi Jiang, Su-Jun Deng, H. Eric Xu

## Abstract

The Omicron variant of SARS-CoV-2 has rapidly become the dominant infective strain and the focus efforts against the ongoing COVID-19 pandemic. Here we report an extensive set of structures of the Omicron spike trimer by its own or in complex with ACE2 and an anti-Omicron antibody. These structures reveal that most Omicron mutations are located on the surface of the spike protein, which confer stronger ACE2 binding by nearly 10 folds but become inactive epitopes resistant to many therapeutic antibodies. Importantly, both RBD and the closed conformation of the Omicron spike trimer are thermodynamically unstable, with the melting temperature of the Omicron RBD decreased by as much as 7°C, making the spiker trimer prone to random open conformations. An unusual RBD-RBD interaction in the ACE2-spike complex unique to Omicron is observed to support the open conformation and ACE2 binding, serving the basis for the higher infectivity of Omicron. A broad-spectrum therapeutic antibody JMB2002, which has completed Phase 1 clinical trial, is found to interact with the same two RBDs to inhibit ACE2 binding, in a mode that is distinguished from all previous antibodies, thus providing the structural basis for the potent inhibition of Omicron by this antibody. Together with biochemical data, our structures provide crucial insights into higher infectivity, antibody evasion and inhibition of Omicron.

---

The Omicron variant of SARS-CoV-2 (B.1.1.529), the causative virus of COVID-19, was initially reported from South Africa on November 24 of 2021, and has become the dominant strain in less than 40 days, with daily infections exceeding one million cases worldwide and three quarters of which are infected with Omicron (*1*). Phylogenetic tree analyses reveal that Omicron is evolved independently from previous variants of concerns (VOC), including the predominant Alpha, Beta, Gamma, and Delta variants (Fig. 1A) (*2-5*). Compared to the original wildtype (WT) strain of SARS-CoV-2, Omicron has an unusually high number of 60 mutations, over 50% of which (37 mutations) are located to the spike protein, the target of most COVID-19 vaccines and therapeutic antibodies (Fig. 1B)(*6-8*). This level of mutations has provided the genetic basis for Omicron to have the astounding ability in transmission, antibody evasion and vaccine resistance. However, the biochemical and structural mechanisms for Omicron’s unusual capability remain unknown and require urgent studies given the astonishing rate of Omicron infections.

**Fig 1.**
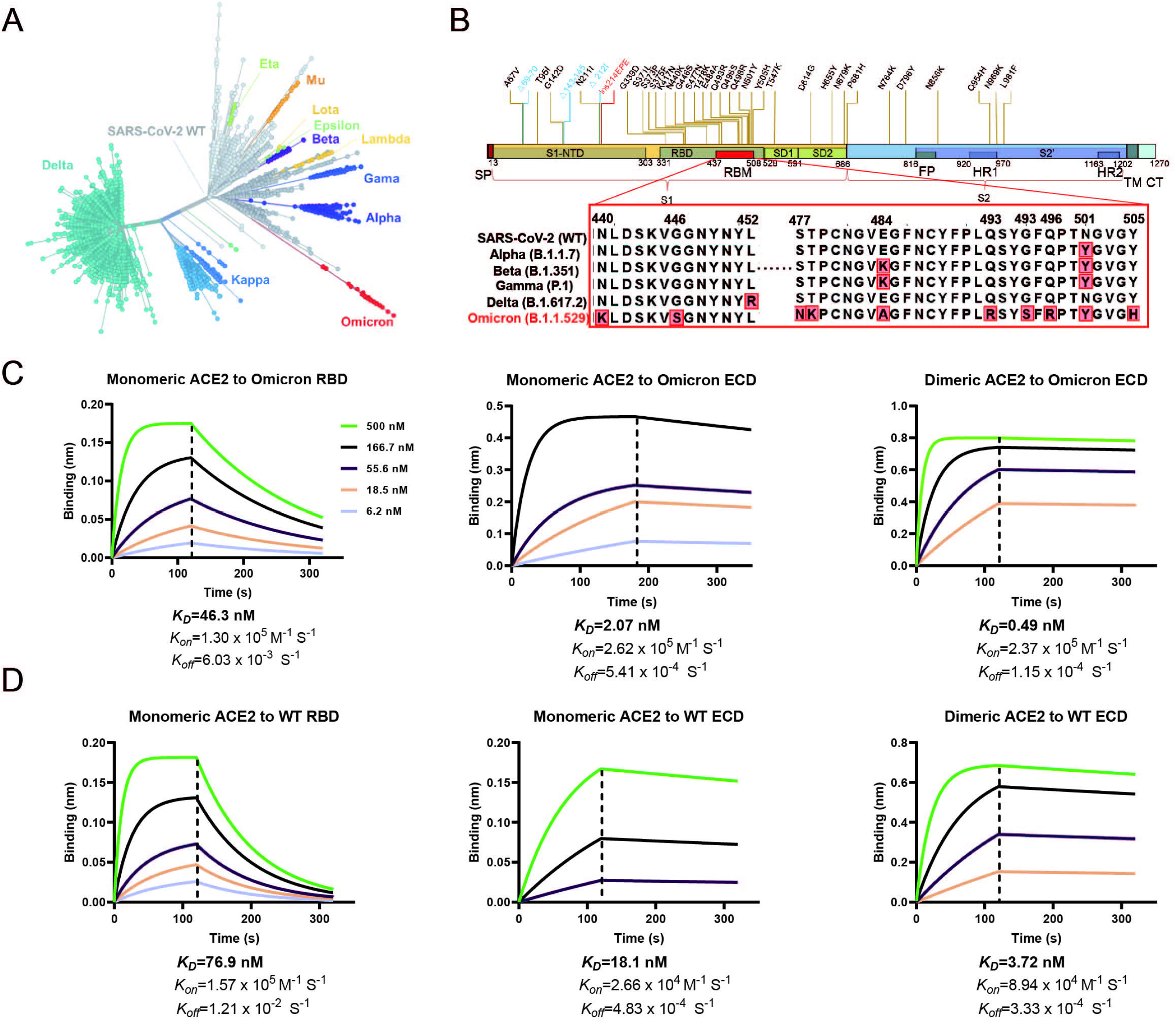
SARS-CoV-2 Omicron spike protein with higher affinity to human ACE2 compared with SARS-CoV-2 wildtype. **(A)** Phylogeny of the SARS-CoV-2 variants. Variants of concern and variants of interest are labelled on the graph, the number of spike protein mutations are positively correlated with the distance from the original strain. The tree is constructed from the Nextstrain servers. **(B)** Schematic of Omicron spike protein domain architecture. The mutations of Omicron spike protein are labeled with different colors (blue for deleting mutation, red for inserting mutation). Mutations in RBM are compared with wildtype SARS-CoV-2 and four other variants of concern strains. SP, signal peptide; RBD, receptor binding domain; RBM, receptor binding motif; SD1, subdomain 1; SD2, subdomain 2; FP, fusion peptide; HR1, heptad repeat 1; HR2, heptad repeat 2; TM, transmembrane region; CT, cytoplasmic tail. **(C-D)** Binding of Omicron spike protein and wildtype SARS-CoV-2 spike protein to human ACE2 determined by BLI. K_D_ values were determined with Octet Data Analysis HT 12.0 software using a 1:1 global fit model.

To study the biochemical mechanism for Omicron’s enhanced transmission, we first biochemically characterized the interactions of the SARS-CoV-2 receptor ACE2 with the spike trimer from Omicron and the original WT strain. The monomeric human ACE2 bound to immobilized Omicron spike protein (K_D_=2.07 nM) with approximately 9-fold higher affinity compared to that of WT spike protein (K_D_=18.1 nM). The dimeric human ACE2 bound to Omicron spike protein (K_D_=0.49 nM) with approximately 8-fold higher affinity compared to that of WT (K_D_=3.72 nM) (Fig. 1C and 1D). We then studied the interactions of ACE2 with isolated RBD from Omicron and WT strains. The monomeric human ACE2 bound to immobilized Omicron RBD (K_D_=46.3 nM) with approximately 1.7-fold higher affinity compared to that of WT RBD (K_D_=76.9 nM) (Fig. 1C and 1D). In biochemical investigation, enhanced interaction of Omicron spike and RBD proteins with human ACE2 was observed. These data were in agreement with previously published data (*9*), demonstrating the higher affinity of Omicron spike and RBD proteins to the SARS-CoV-2 receptor ACE2, thus conferring the increased infectivity of the Omicron variant.

To determine the structural basis of higher affinity of the Omicron spike protein to ACE2, we solved the structure of the ACE2-Omicron spike protein complex at a global resolution of 2.77 Å by using a purified ACE2-spike complex (Fig. S1A). Despite the excess amount of ACE2 (molar ratio of 3.2 ACE2 to 1 spike trimer), we only observed strong density for one ACE2 bound to one RBD from the spike trimer in the open “up” conformation (Fig. 2A, and S2), where the other two RBDs, with clear density, are in close “down” conformation. Surprisingly, particle classification revealed that the majority of picked particles (~70%) are absence of ACE2. The overall structure of the apo Omicron spike trimer reached a global resolution at 2.56 Å and resembles the structures of original WT strain (*10-12*) and previous variants of concern (*13, 14*). In contrast to the clear visibility of the three RBDs in the ACE2-Omicron spike complex, the three RBDs in the apo Omicron spike trimer are nearly invisible in high resolution maps (Fig. S3A), and low resolution maps reveal that all RBDs are visible in down conformation (Fig. S3B), indicating that the RBD in the apo Omicron Spike complex is highly dynamic, preferentially in down state, whereas ACE2 binding helps stabilize the conformation of the three RBDs.

**Fig 2.**
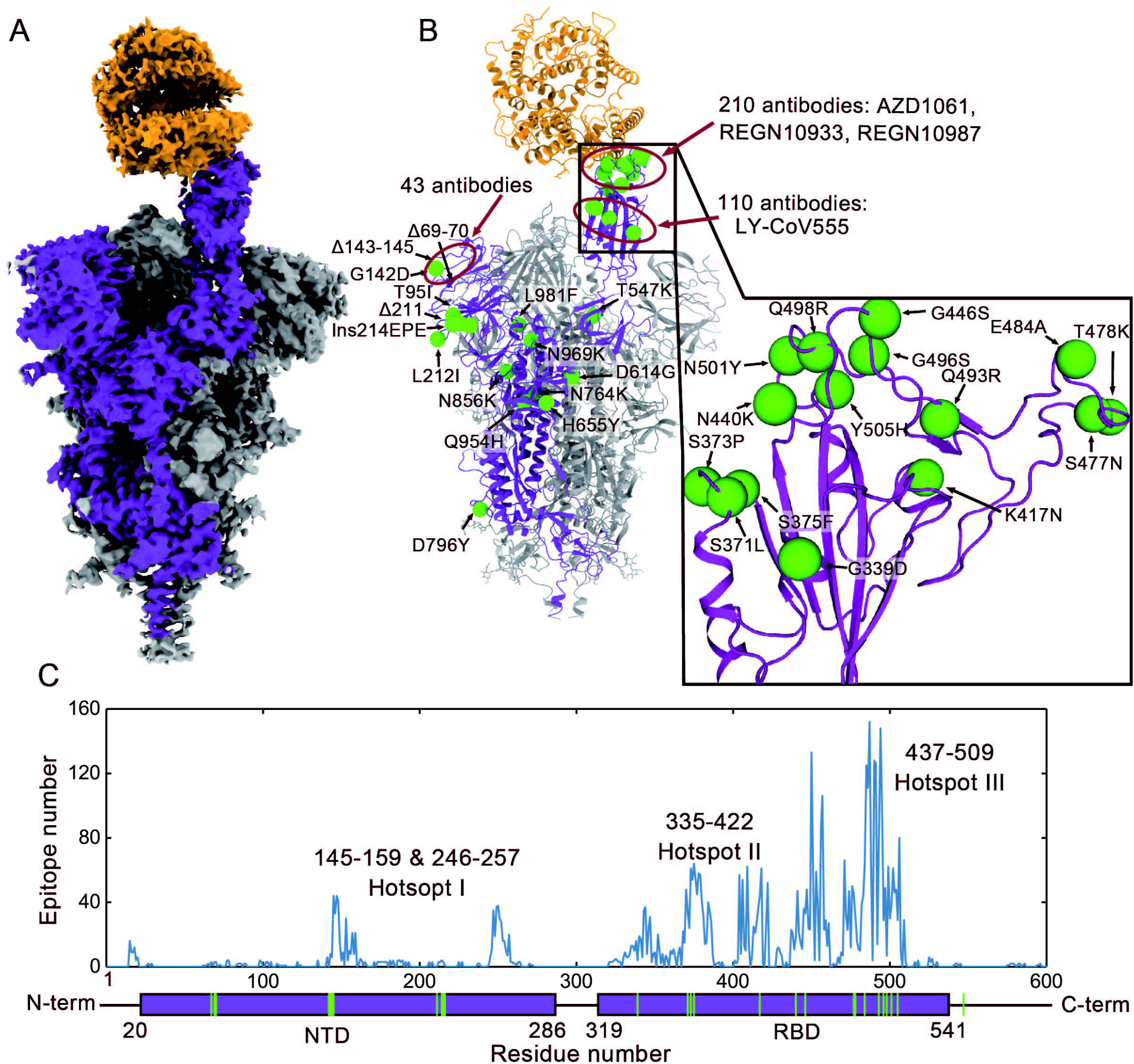
The structure of ACE2 Omicron spike trimer complex and epitopes of current antibodies. **(A)** Cryo-EM density of the ACE2-Omicron spike trimer complex. **(B)** The overall structure of the ACE2-Omicron spiker trimer complex and the locations of Omicron mutations. Epitope hotspots are highlighted in red circles with the number of antibodies indicated next to to the epitopes. **(C)** The histogram of epitope corresponding with residue number. Each epitope was counted if more than 3 heavy atoms of this residue are closer than 5 Å with the antibody. The structural component of SARS-CoV-2 is shown below.

To examine the dynamic nature of the Omicron RBD, we performed thermal shift assays, which revealed that both the Omicron and WT spike trimer displayed two melting temperatures with identical TM2 of 71.5 ° C for both spike trimers but different TM1 by as much as 7.1 °C (Fig. S1C), which is nearly identical to the Tm difference between the isolated Omicron and WT RBDs (Fig. S1D), suggesting that the Omicron RBD is much less stable than the WT RBD in the spike trimer form or in isolated monomeric form (Fig. S3). We further confirmed the highly flexible nature of the Omicron RBD by performing hydrogen-deuterium exchange mass spectrometry (HDX), which showed that the Omicron spike trimer has overall higher rate of HDX (Fig. S4), particularly in the RBD region, consistent with its lower thermal stability. The instability of the Omicron RBD would probably synergizes with its higher binding affinity with ACE2 in promoting Omicron infectivity.

Mapping of the 37 mutations onto the up protomer of the ACE2 bound spike trimer reveals that most mutations are located on the surface of the spike protein, with many of them serving as the epitopes of therapeutic antibodies (Fig. 2B). Eight mutations in the NTD would affect the structures of the epitopes for a number of antibodies, for example, Δ143-145 would remove the epitope for 4A8 antibody (*15*). 15 mutations are in RBD, which contains the sites for ACE2 binding as well as the site for 90% of antibodies induced by infection or vaccination, are clustered into two hot spots (II and III), with 10 mutations in RBM and 5 mutations near the core structure domain (Fig. 2C). Both hot spots are the major epitopes for therapeutic antibodies LY-CoV 555, AZD1061, REGN10987. (*16-18*)

Local refinement of the ACE2-RBD region produced a high-quality map at 2.57 Å resolution, which allowed unambiguous building of the ACE2-RBD complex (Fig. 3A, Table S1). To our surprise, the overall structure of ACE2-RBDOmi complex is exceedingly similar to the two high resolution X-ray structures of the ACE2-RBDWT complex (PDB codes: 6LZG and 6M0J)(*19, 20*), with the Cα atoms of the whole RBD deviated less than 0.4 Å (Fig. 3B, Table S2), despite the initial model used in our structure was from a cryo-EM structure (PDB code: 7A94)(*21*). Even more surprisingly, the backbone structure of the Omicron RBM is nearly identical to the WT RBM with RMSD of its Cα atoms less than 0.35 Å despite that there 10 mutations out of this 72 residues region. The extreme similarity of our ACE2-RBD^Omi^ complex structure to the two independent high resolution X-ray structures of the ACE2-RBD^WT^ complex reflects the high accuracy and quality of our cryo EM maps and structure modeling. The major differences residues in the ACE2-RBM interface, in which the Omicron RBM forms extra interactions with ACE2, including interactions from RRM mutations S477N, Q493R, Q496S, Q498R, and N501Y to ACE2 (Fig. 3C). In particular, the side chain of S477N forms two extra hydrogen bonds with S19 of ACE2; the Q498R mutation from two additional hydrogen bonds with Q42 and D38 from ACE2; and the N501Y mutation forms extensive packing interactions with ACE2 residues Y41 and K353. These additional interactions may compensate for the loss interactions caused by RBD mutations K417N, E484A and Y505H, which WT RBD residues forms polar interactions with ACE2 (Fig. 3C) (*22*).

**Fig 3.**
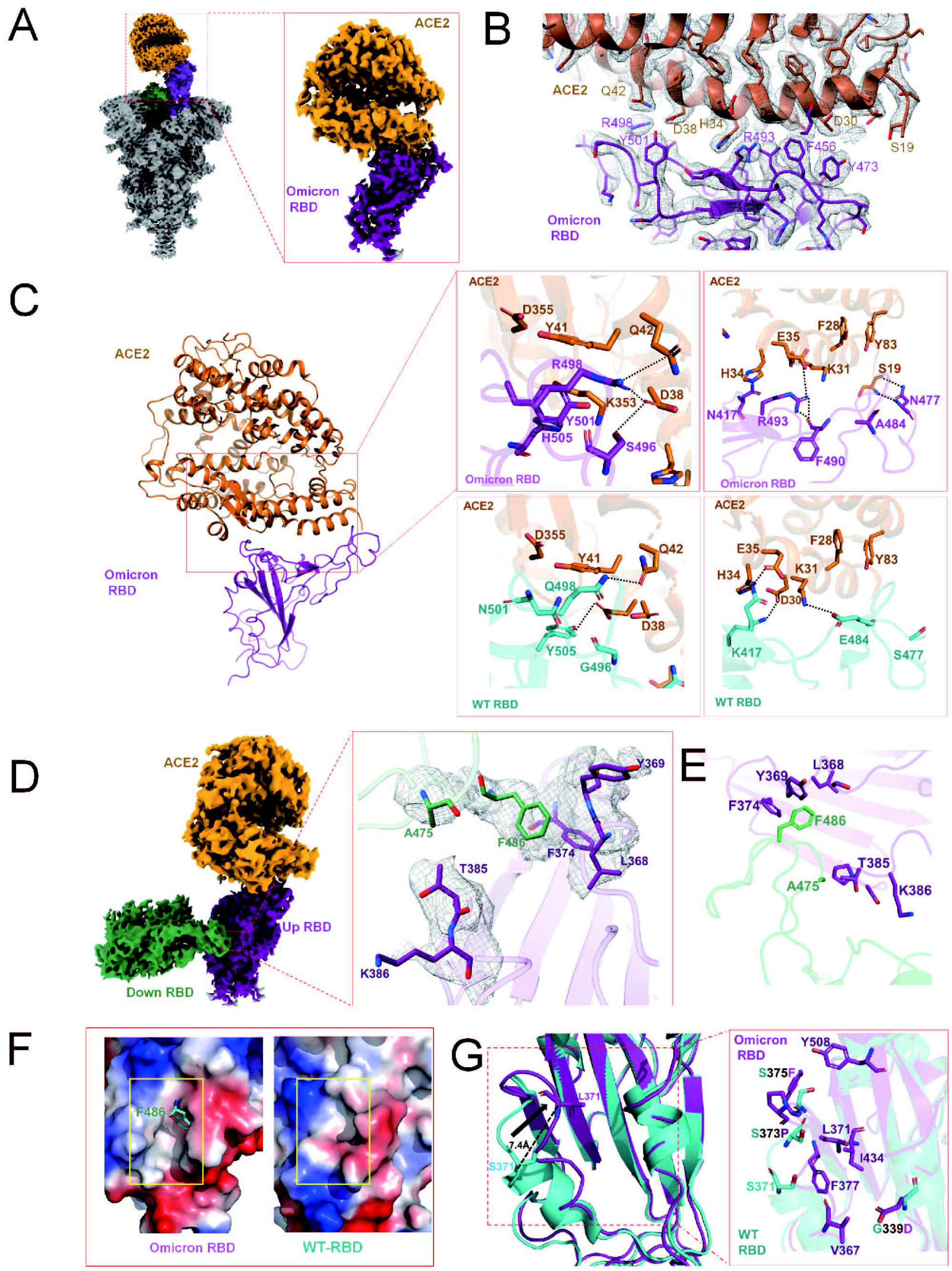
Structural analysis of Omicron RBD and ACE2. **(A)** Cryo-EM density and location of Omicron RBD-ACE2 bound region. ACE2 is colored in orange. The Omicron ECD-trimer is colored in grey with two RBDs colored in purple and green. **(B)** ACE2-RBD binding interface. ACE2 is colored in orange and the RBD is colored in purple. Residues are shown in sticks with the correspondent cryo-EM density represented in mesh. **(C)** Comparison of Omicron RBD-ACE2 and WT RBD-ACE2 interfaces. Left panel, an overall structural model of Omicron RBD-ACE2 bound region. Right-up panel, zoomed-in view of Omicron RBD-ACE2 with hydrogen bonds interactions. Right-down panel, the detailed hydrogen bonds interactions in WT RBD-ACE2 interfaces with the same view as in right-up panel. WT RBD is sea blue, Omicron RBD is purple, and ACE2 is orange. Hydrogen bonds interactions are in dotted lines. **(D)** The interaction interface between down RBD and ACE2-RBD. Left panel, overall cryo-EM density of down RBD-ACE2-RBD region. Right panel, a close-up view of RBD-RBD interaction. ACE2 bound RBD, also named up RBD here is in purple. The down RBD, which directly bounds to up RBD is in green. **(E)** The contribution of F486 and A475 from the down RBD in RBD-RBD interactions. F486 and A475 from down RBD in green, and F374, Y369, L368, T385, and K386 from up RBD in purple are shown in sticks. **(F)** Comparison of electrostatic potential maps between Omicron RBD (left panel) and WT RBD (right panel). Down RBD bound region in Omicron RBD and the correspondent region in WT RBD are highlighted in yellow boxes. Positive and negative charged regions are in blue and red, respectively. **(G)** The mutation-induced conformations change of Omicron RBD comparing with WT RBD.

Another surprising observation is an RBD-RBD interactions from one of the two down RBDs to the up RBD, with the interface comprised by residues A475, F486, and Y489 from the down RBD and residues L368, Y369, F374, T385 and K386 from the up RBD (Fig. 3D, 3E). Structure comparison reveals that the RBD-RBD interface is not observed within the WT spike trimer because of a large movement of a loop (residues 368-374) as large as 7.4 Å, caused by Omicron RBD mutations S371L, S373P, G339D, and S375F, which also comprise a hotspot of epitopes (Fig. 3F, 3G, and Fig. 2C). This new RBD-RBD interaction may stabilize the up conformation of RBD that promotes ACE2 binding. Consistently, ACE2 binds to the Omicron spike trimer with 8-9 folds higher affinity than to the WT spiker trimer, but only 1.7 fold stronger to the RBD monomer (Fig. 1C, 1D). Together, our biochemical data with these structural observations provide a mechanistic basis for the higher infectivity of the Omicron variant than the WT strain.

We had previously discovered an antibody, JMB2002, which showed potent efficacy against the WT SARS-CoV-2 in cell-based model as well as in rhesus monkey model (*23*). JMB2002 has finished Phase I clinical trial in healthy donors with excellent safety and PK properties and has been approved for clinical trial in US (IND 154745). To evaluate the neutralizing activity of JMB2002 against the Omicron variant, the binding affinity of JMB2002 to WT and Omicron spike proteins was determined. JMB2002 Fab recognized Omicron spike protein with 4-fold increased binding affinity (K_D_=5.32 nM) compared to WT spike protein (K_D_=20.4 nM). JMB2002 bound to the Omicron spike trimer with slightly higher affinity (K_D_=0.47 nM) in comparison to that of WT (K_D_=0.63 nM) (Fig. 4A-4D). Furthermore, JMB2002 was able to directly inhibit the binding of ACE2 to the Omicron spike trimer with an IC_50_ of 1.8 nM (Fig. 4E). In pseudovirus neutralization assays, JMB2002 effectively blocked the entry of Omicron pseudovirus into human ACE2 expressing cells in addition to its blocking of the WT pseudovirus (Fig. 4F and Fig. S5). JMB2002 was also able to neutralize a number of VOCs, including variants of Alpha, Beta, and Gamma, but not Delta (Fig. S5). Thus, JMB2002 may be an effective therapy against all WHO VOC except the Delta variant.

**Fig 4.**
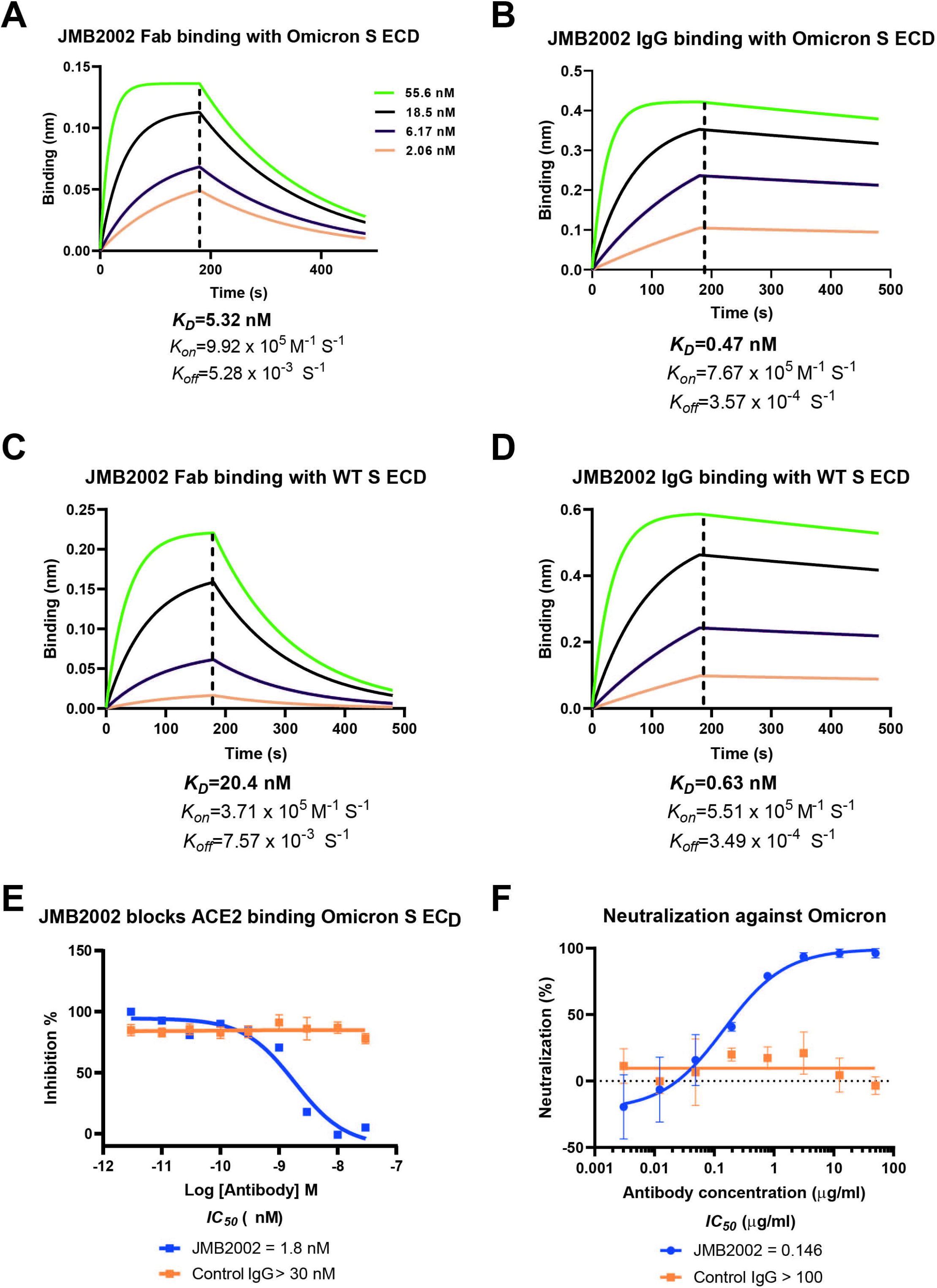
Inhibition of ACE2 binding to the spike trimer by an anti-Omicron antibody. **(A and C)** Binding of JMB2002 Fab to Omicron and WT spiker trimer. **(B and D)** Binding of JMB2002 IgG to Omicron and WT spiker trimer. **(E)** Direct inhibition of ACE2 binding to the Omicron spike trimer by JMB2002. **(F)** Inhibition of the pseudo virus of WT and omicron by JMB2002.

To reveal the basis of JMB2002 inhibition of Omicron, we solved the structure of the Omicron spike trimer bound to a Fab from JMB2002 at a global resolution of 2.69 Å (Fig. 5A, S1B, S5, and Table S1). To stabilize the constant regions of Fab, we used a nanobody that binds to the interface between the variable and constant regions of the heavy chain of Fab (*24*). The EM density maps reveal clear arrangement of the Fab-spike trimer complex, with the binding of 2 Fab molecules to the same two RBDs (one up and one down) from the ACE2-spike complex (Fig. 5A-5G). The overall structure of the spike trimer from the Fab-bound complex is very similar to that of the ACE2-bound complex, with an RMSD of 1.0 Å for the entire Cα atoms of the spiker trimer, including the RBD-RBD dimer configuration that is unique to Omicron (Fig. S7A and S7B).

**Fig 5.**
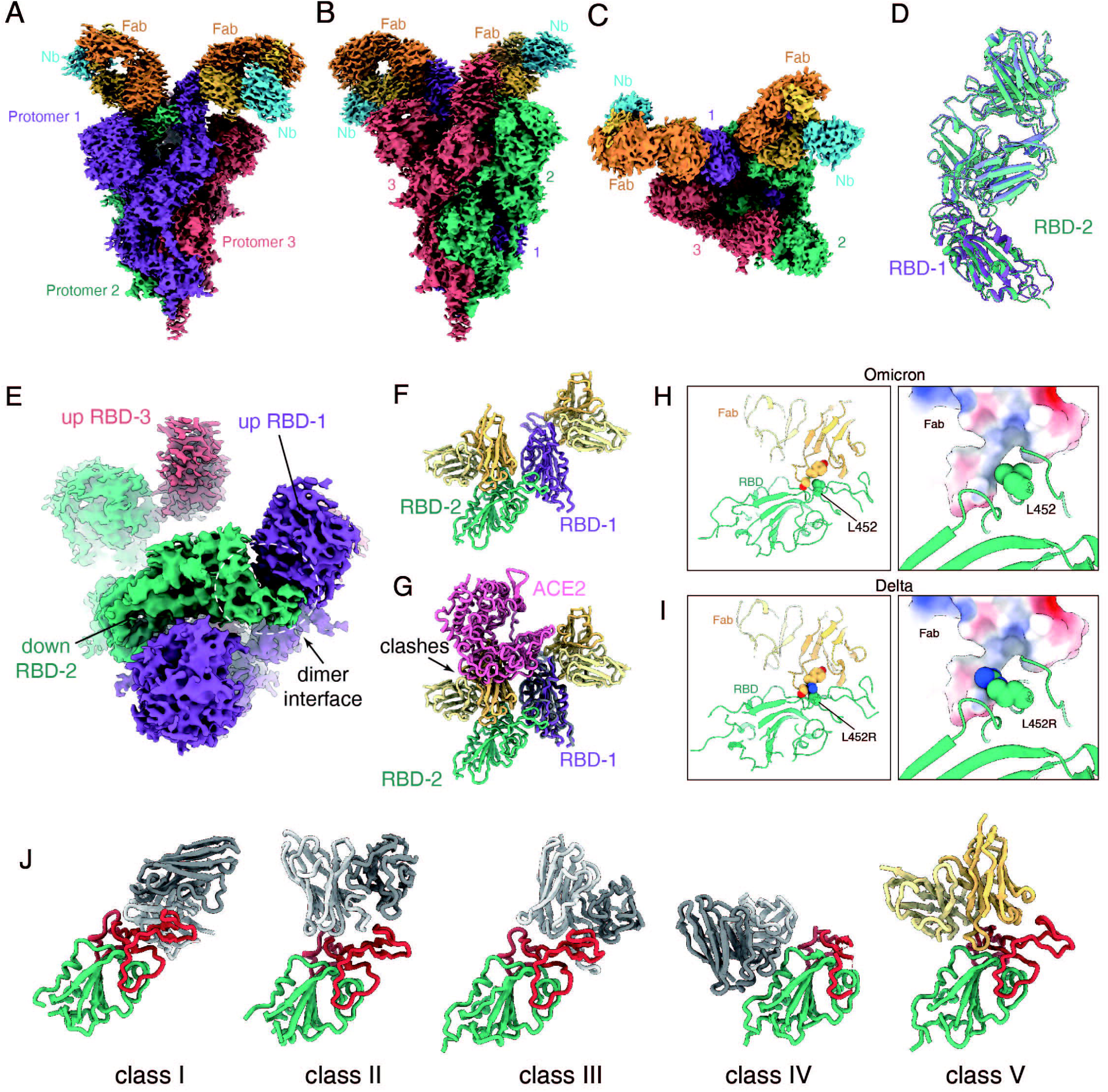
Structure of Omicron S ECD antibody complex. **(A-C)** Cryo-EM density map of the Fab-bound Omicron S protein. The map showed with two front views **(A and B)** and a top view **(C). (D)** Superposition of the RBD-1 and RBD-2 both bound to a Fab molecule. **(E)** The interface between the up state RBD-1 and the down state RBD-2 of Omicron Spike protein. **(F)** The structure of the RBD-1 and RBD-2 bound to Fab. **(G)** Superposition of the ACE2-bound and Fab-bound RBD-1 shows the binding of another Fab to RBD-2 inhibits ACE2 binds to RBD-1. **(H)** Omicron variant RBD L452 residue formed interactions with Fab. **(I)** The Delta variant L452R mutation clashes the Fab binding. **(J)** Binding modes of 4 representative 4 classes of antibody that neutralize SARS-CoV-2. PDB codes: class I, 7CM4; class II, 7CHF; class III, 7K90; class IV, 6WPS. The Fab in the Omicron S protein structure shows distinct binding modes from other 4 classes of antibodies.

Within the Fab-spike trimer structure, both Fab binds to the same region in the RBD (Fig. 5D). Within the isolated RBD, the Fab binding site does not overlap with the ACE2 binding site (Fig. S7C). However, within the spike trimer, Fab binding to the down RBD will clash with ACE2 binding to the up RBD (Fig. 5G), preventing the ACE2 binding in a conformation-specific state. Thus, the JMB2002 Fab inhibits ACE2 binding to the spike trimer through stabilizing a specific conformation that is clashed with ACE2 binding. Simultaneous binding of two Fab molecules to the spike trimer is consistent with 10-fold higher affinity of the IgG format of JMB2002 than the Fab format (Fig. 5A and 4A-4B).

Particle classification also revealed two additional antibody-bound complexes at a global resolution of 2.94 Å and 3.18 Å, respectively (Fig. S5 and S8). One of the two complexes has the spike trimer with two-up RBDs and one down RBD, with each RBD bound to one Fab (Fig. S8A). The other complex contains the spike trimer with one up-RBD bound to one Fab and two-down RBDs in the apo state (Fig. S8B). The diverse configuration and stoichiometry ratio of the spike trimer bound to the antibody further highlight the conformation flexibility of the Omicron spike RBD. The ability of the spike trimer bound to three Fab molecules also provide additional basis for the blocking ability of JMB2002 to Omicron.

The L452R mutation in the Delta variant is at the center of the binding epitope of JMB2002 and this mutation will clash with Y102 from the heavy chain of Fab, thus providing an explanation for its loss of potency to the Delta variant (Fig. 5H, 5I). In addition, the binding site of the JMB2002 Fab to the RBD site in the spike trimer which is distinct from all previous class I to class IV antibodies (Fig. 5J), thus JMB2002 represents a new class of antibody to the spike trimer.

The astonishing spread of Omicron infections has taken the whole World to a great surprise, with extensive medical resources devoted to overcome this new wave of pandemic(*25*). Mechanistic studies into the rapid pace of infection and immune evasion comprise of an integrated effort to combat Omicron infections. In this paper, we report three structures of the Omicron spike protein by its own or bound to the SARS-CoV-2 receptor ACE2 and a broad-spectrum therapeutic antibody. Combined with the extensive biochemical and biophysical data, our results display that the extensive mutations in the Omicron spike proteins not only destabilize the closed conformation of the three RBD within the spike trimer but also increase the direct binding affinity by nearly 10 folds to its receptor ACE2, both of these properties serve as the basis for the higher infectivity of Omicron. The structure of the ACE2 bound Omicron spiker trimer reveal an unusual RBD-RBD interaction unique to Omicron and extra interactions in the ACE2-RBD interface, both of which contribute to the higher affinity of ACE2 to the Omicron spiker trimer, which could therefore lead to the higher infectivity of Omicron. Structural analyses of the Omicron spike trimer also reveal that its extensive mutations are mostly located on the hotspots of epitopes, providing a mechanistic basis for the outstanding capability of Omicron to escape most therapeutic antibodies and vaccinations.

In addition to revealing the mechanisms of increased infectivity and immune evasion, our antibody-bound Omicron spiker trimer also uncovers a distinguished mode of antibody binding to the spike trimer, in which the unique RBD-RBD configuration is preserved. The binding epitope of this broad-spectrum antibody is different from all previous anti-SARS-CoV-2 antibodies, therefore opening a new venue for antibody drug discovery targeting various strains of SARS-CoV-2, including Omicron.

## Supporting information

full text

The PDB for the Omicron spike RBD bound to ACE2

