## Supplementary material for "Structures of the Omicron spike trimer with ACE2 and an anti-Omicron antibody: mechanisms for the high infectivity, immune evasion and antibody drug discovery": full text

**Supplementary Information**

#### Materials and Methods

##### Protein expression and purification

The N-terminal peptidase domain of human ACE2 (residues Ser19–Asp615) with an N-terminal GP67 signal peptide for secretion and a C-terminal 8×His tag for purification was inserted into a modified pFastBac vector (Invitrogen). All constructs were generated using the Phanta Max Super-Fidelity DNA Polymerase (Vazyme Biotech Co.,Ltd) and verified by DNA sequencing (Genewiz). Hi5 cell cultures were grown in ESF 921 serum-free medium (Expression Systems) to a density of 2-3 million cells per ml and then infected with baculoviruses for human ACE2 at a multiplicity of infection (m.o.i.) of about 5. The supernatant of cell culture containing the secreted peptidase domain of human ACE2 was collected 60 h after infection, and then bound to Ni-NTA beads (Smart-Lifesciences). After wash with buffer without imidazole, the protein was eluted with buffer containing 300 mM imidazole, concentrated and then size-separated by a Superdex 200 Increase10/300 GL column in 25 mM HEPES pH 7.4, 150 mM sodium chloride. The fractions for the peptidase domain of human ACE2 were collected, concentrated to about 2 mg/ml, and stored at − 80 °C until use.

The SARS-CoV-2 Spike ECD and RBD proteins were all purchased from Sino Biological Inc., including Omicron spike ECD protein, Omicron spike RBD protein, WT spike ECD protein, WT spike RBD protein, Delta spike ECD protein, Delta spike RBD protein.

Nanobody (Nb) with a C-terminal His6 tag was expressed in the periplasm of *E.coli* strain BL21(DE3) bacteria (NEB). Cultures of 2 L cells were grown to OD600 = 0.8 at 37 °C, 180 r.p.m. in 2×YT media containing 100 μg/mL ampicillin. Then 0.1mM IPTG was added to the medium to induce protein expression at 28°C, 180 r.p.m. for another 8 h. Cells were harvested by centrifugation (5316 g, 30 min) and disturbed in ice-cold buffer (20 mM HEPES pH 7.4, 100 mM NaCl), then centrifuged to remove cell debris. Nb was purified by nickel affinity chromatography as previously described followed by size-exclusion chromatography using a HiLoad 16/600 Superdex 75 column. Selected fractions of Nb were finally concentrated to ~2 mg/ mL with 10% glycerol and kept frozen at -80 °C for further use. The quality of purified proteins was assessed by SDS-PAGE.

**Protein complex formation**

For the ACE2 bound Omicron spike ECD protein complex, the Omicron spike ECD protein was incubated with purified peptidase domain of human ACE2 at molar ratios of 1:3.2 (spike trimer to peptidase domain of human ACE2) for 60 min on ice prior to purification by gel filtration chromatography using a Superose6 column (GE Healthcare) pre-equilibrated with TBS buffer (20 mM Tris, pH8.0, 150 mM NaCl). For the JMB2002Fab bound Omicron spike ECD protein complex, the Omicron spike ECD protein was incubated with JMB2002Fab d anti-Fab nanobody at molar ratios of 1:4:5 (spike trimer to JMB2002Fab to anti-Fab nanobody) for 60 min on ice prior to purification by gel filtration chromatography using a Superose6 column (GE Healthcare) pre-equilibrated with HBS buffer (25 mM HEPES, pH 7.5, 150 mM NaCl). For the Omicron spike ECD trimer protein only, the Omicron spike ECD protein was directedly set to purification by gel filtration chromatography using a Superose6 column (GE Healthcare) pre-equilibrated with TBS buffer (20 mM Tris, pH 8.0, 150 mM NaCl) or HBS buffer (25 mM HEPES, pH 7.5, 150 mM NaCl). Fractions containing the complex were pooled and concentrated to 1.5-3 mg/ml.

**Cryo-EM sample preparation and data acquisition**

For the Omicron spike ECD trimer protein only at pH 8.0, 3 μL of the Omicron spike ECD purified with TBS buffer at the concentration of 1.9 mg/ml were applied to glow-discharged UltrAuFoil Holey Gold 300 mesh grids (Au, R0.6/1); For the Omicron spike ECD trimer protein only at pH 4.0, the Omicron spike ECD purified with HBS buffer at the concentration of 1.8 mg/ml was mixed with 1 M sodium acetate, pH4.0

(final sodium acetate concentration: 0.1 M), then 3 μL sample were applied to glow-discharged UltrAuFoil Holey Gold 300 mesh grids (Au, R0.6/1); For the ACE2 bound Omicron spike ECD protein complex, 3 μL sample purified with TBS buffer at the concentration of 2.5 mg/ml were applied to glow-discharged UltrAuFoil Holey Gold 300 mesh grids (Au, R1.2/1.3); For the JMB2002Fab bound Omicron spike ECD protein complex, 3 μL sample purified with HBS buffer at the concentration of 2.5 mg/ml were applied to glow-discharged 300 mesh amorphous NiTi foil (Au, 1.2/1.3). Extra samples were blotted with filter paper for 3.0 s and plunge-frozen in liquid ethane using a FEI Vitrobot Mark IV. Cryo-EM micrographs were collected on a 300kV Titan Krios microscope (FEI) equipped with a K3 direct detection camera. For ECD-ACE2 dataset, 8,194 movies were collected, and for ECD-Fab dataset, 14,740 movies were collected on a Titan Krios equipped with a Gatan K3 direct electron detection device at 300kV with a magnification of 105,000, corresponding to a pixel size 0.824 Å. Image acquisition was performed with EPU Software (FEI Eindhoven, Netherlands). We collected a total of 36 frames accumulating to a total dose of 50 $e^{-}$ $Å^{-2}$ over 2.5 s exposure.

**Cryo-EM data processing**

For ECD-ACE2 dataset, MotionCor2 was used to perform the frame-based motion-correction algorithm to generate drift-corrected micrograph for further processing and CTFFIND4 provided the estimation of the contrast transfer function (CTF) parameters(*1, 2*). 1,566 aligned micrographs were deleted because of contaminations or bad ice quality. After selection, approximately 1,500 particles were manually picked and two-dimensional (2D) classes were calculated and used as references for automatic picking. A total of 5,071,668 particles were extracted from the cryo-EM micrographs performed with Relion3.0(*3*). All subsequent steps including 2D classification, three-dimensional (3D) classification, 3D refinement, and local refinement were performed using cryoSPARC(*4*), unless stated otherwise. Two rounds of reference-free 2D classification, yielding 854,875 particles after clearance.

Three rounds hetero refinement separated out 522,186 particles that resulted to a clearer density of ECD-trimer structure at 2.56 Å global resolution. Two rounds hetero refinement separated out 242,374 particles that resulted to a clearer density of ACE2 bound ECD-trimer structure at 2.77 Å global resolution. Local refinement focused on the ACE2-RBD or RBD-ACE2-RBD with masks could reconstitute ACE2-RBD and RBD-ACE2-RBD structures at 2.79 Å and 2.57 Å global resolution, respectively. Local resolution estimate was performed with cryoSPARC.

For ECD-Fab dataset, MotionCor2 was used to perform the frame-based motion-correction algorithm to generate drift-corrected micrograph for further processing and CTFFIND4 provided the estimation of the contrast transfer function (CTF) parameters (*1, 2*). Automated particle selection yielded 6,563,376 particles. Two rounds of reference-free 2D classification and three rounds hetero refinement separated out 172,001 particles that resulted to a density of ECD-trimer bound to 1 Fab at 3.18 Å global resolution, 906,588 particles that resulted a density of ECD-trimer bound to 2 Fab at 2.69 Å global resolution, and 434,893 particles that resulted to a density of ECD-trimer bound to 1 Fab at 2.92 Å global resolution. The maps were sharpened by DeepEMhancer (*5*). Local resolution estimate was performed with cryoSPARC.

**Model building**

ECD structure from Swiss-model prediction was used as the starting reference model for ECD-trimer building(*6*). Structure of ACE2 derived from PDB entry 7KMB(*7*) was rigid body fit into the density. For the Fab-ECD complex, the structure of the SARS-CoV-2 spike glycoprotein (PDB: 6VXX(*8*)) and the Fab-nanobody-AMPK structure (PDB:7JHH(*9*)) were used as initial model. All models were fitted into the EM density map using UCSF Chimera(*10*) followed by iterative rounds of manual adjustment and automated rebuilding in COOT(*11*) and PHENIX(*12*), respectively. The model was finalized by rebuilding in ISOLDE(*13*) followed by refinement in PHENIX with torsion-angle restraints to the input model. The final model statistics were validated using

Comprehensive validation (cryo-EM) in PHENIX(*12*). All structural figures were prepared using Chimera(*10*), Chimera X(*14*), and PyMOL (Schrödinger, LLC.).

**Thermal shift assay (TSA)**

The spike ECD and RBD proteins of Omicron and WT SARS-CoV-2 were diluted to 0.5 mg/ml using 200 mM HEPES pH 7.4, 150 mM sodium chloride (final HEPES concentration: 100 mM). Then the diluted proteins were mixed with 10× SYPRO Orange (Thermo Fisher) and incubated for 10 min at room temperature. The reaction was performed in 96-well plates with a final volume of 20 μl. The thermal melting curve were monitored using a LightCycler 480 II Real-Time PCR System (Roche Diagnostics) with a ramp rate of 3.6 °C per minute from 25 °C to 80 °C. The melting peaks were calculated by the LightCycler 480 software provided by Roche Diagnostics.

**Measurement of human ACE2 binding to Omicron and WT spike RBD/ECD by biolayer interferometry**

The binding of human ACE2 to Omicron and WT spike RBD/ECD were performed using Octet Red96e (Sartorius). The biotinylation of SARS-CoV-2 spike RBD or monomeric human ACE2 protein were carried out by using EZ-Link Sulfo-NHS-LC-Biotin kit (ThermoFisher Scientific, A39257) following the manufacturer's instruction. SA biosensors (Sartorius, 18-5020) were used to immobilize the biotinylated WT spike RBD protein (Sino Biological, 40589-V08H), Omicron SARS-CoV-2 spike RBD protein (Sino Biological, 40952-V08H121) or monomeric human ACE2 protein. The association of SARS-CoV-2 spike RBD-coated sensors with different concentrations of monomeric human ACE2 were recorded prior to 10 min dissociation in kinetics buffer (0.02% Tween-20 in PBS). The binding of biotinylated monomeric human ACE2 to WT SARS-CoV-2 spike ECD (Sino Biological, 40589-V08B1) or Omicron SARS-CoV-2 spike ECD (Sino Biological, 40589-V08H26) was also evaluated. Protein A biosensors (Sartorius, 18-5012) were used to capture the dimeric human

ACE2 (Sino Biological, 10108-H02H) followed by measuring the association and dissociation with WT or Omicron spike ECD protein. $K_D$ values were calculated with Octet Data Analysis HT 12.0 software using a 1:1 global fit model. Data were plotted using Prism V8.0 software (GraphPad).

**Measurement of anti-SARS-CoV-2 antibody JMB2002 binding to Omicron and WT spike ECD by biolayer interferometry**

The interaction of anti-SARS-CoV-2 antibody JMB2002 with Omicron and WT spike ECD was determined using Octet Red96e (Sartorius). SA biosensors (Sartorius, 18-5020) were used to capture the biotinylated WT SARS-CoV-2 spike ECD protein (ACRO, SPN-C82E9) or Omicron SARS-CoV-2 spike ECD protein (ACRO, SPN-C82Ee). The association of SARS-CoV-2 spike protein-coated sensors with different concentrations of anti-SARS-CoV-2 antibody JMB2002 were recorded prior to 10 min dissociation in kinetics buffer (0.02% Tween-20 in PBS). $K_D$ values were calculated with Octet Data Analysis HT 12.0 software using a 1:1 global fit model. Data were plotted using Prism V8.0 software (GraphPad).

**Pseudovirus neutralization assay**

In the pseudovirus neutralization assay, serial dilutions of JMB2002 IgG were preincubated with an equal volume of WT, Alpha, Beta, Gamma, Delta and Omicron pseudovirus ($2\times10^4$ $TCID_{50}$/mL) for 1 h at 37°C. Subsequently, HEK293 cells stably expressing hACE2 (Vazyme, DD1401) ($2\times10^4$ cells/well) were seeded in 96-well plates, treated with the corresponding pseudovirus and antibody mixtures, and incubated at 37°C for 48 h. Luciferase activity was measured using Bio-Lite Luciferase Assay System (Vazyme, DD1201). The neutralization inhibition rate was calculated with the following formula.

$$\text{Inhibition rate (\%)} = \left(1 - \frac{\text{mean intensity of sample} - \text{mean intensity of blank control}}{\text{mean intensity of negative control} - \text{mean intensity of blank control}}\right) X\,100\%$$

$IC_{50}$ values were calculated by a four-parameter logistic curve fitting approach in Prism V8.0 software (GraphPad).

**AlphaScreen assays for JMB2002 inhibition of ACE2 binding to the Omicron spike ECD trimer**

The JMB2002 ability to block ACE2 binding Omicron S ECD was assessed by luminescence based AlphaScreen technology (PerkinElmer) using a hexahistidine detection kit (*15*). 10 nM ACE2 were attached to streptavidin-coated donor beads, and 10 nM His6-tagged Omicron S ECD were attached to nickel-chelated acceptor beads. Donor beads contain a photosensitizer that, upon activation at 680 nm, converts ambient oxygen into singlet oxygen. When the acceptor beads are brought into close proximity of the donor beads by ACE2-Omicron S ECD interaction, energy is transferred from singlet oxygen to thioxene derivatives in the acceptor beads, resulting in light emission at 520 to 620 nm. The binding mixtures, containing the indicated amounts of ACE2 and Omicron S ECD and streptavidin-coated donor beads (5mg/ml) and Ni-chelate-coated acceptor beads, were incubated in 50 mM Mops (pH 7.4), 100 mM NaCl, and bovine serum albumin (0.1 mg/ml) for 1 to 2 hours before data collection using an EnVision plate reader (PerkinElmer). For the JMB2002 inhibition, increasing concentrations of JMB2002 or Control IgG were added in addition to the ACE2-Omicron S ECD. The $IC_{50}$ values were derived from curve fitting on the basis of a competitive inhibitor model with GraphPad Prism, using conditions where the concentrations of tagged binding partners were below $K_D$.

**HDX detected by mass spectrometry**

HDX-MS were performed as following: 5 µM purified WT or Omicron Spike proteins were prepared for HDX reactions at 4 °C. Four-microliter of protein was diluted into 16 µl D2O on exchange buffer (50 mM HEPES, pH 7.5, 50 mM NaCl, 2 mM DTT) and incubated for various HDX time points (e.g., 0, 10, 60, 300 s) at 4 °C and quenched by mixing with 20 µl of ice-cold 3 M gHCL, 1% trifluoroacetic acid. Each quenched sample was immediately injected into the LEAP Pal 3.0 HDX platform. Upon injection, samples were passed through an immobilized pepsin column (2mm ×

2cm) at 120 µl min$^{-1}$ and the digested peptides were captured on a C18 PepMap300 trap column (ThermoFisher) and desalted. Peptides were separated across a 2.1mm × 5cm $C_{18}$ separating column (1.9 µm Hypersil Gold, ThermoFisher) with a linear gradient of 4% - 40% $CH_3CN$ and 0.3% formic acid, over 6 min. Sample handling, protein digestion and peptide separation were conducted at 4°C.

Mass spectrometric data were acquired using a Fusion Orbitrap mass spectrometer (ThermoFisher) with a measured resolving power of 65,000 at *m*/*z* 400. HDX analyses were performed in triplicate, with single preparations of each protein ligand complex. The intensity weighted mean m/z centroid value of each peptide envelope was calculated and subsequently converted into a percentage of deuterium incorporation. Statistical significance for the differential HDX data is determined by an unpaired t-test for each time point, a procedure that is integrated into the HDX Workbench software. Corrections for back-exchange were made on the basis of an estimated 70% deuterium recovery, and accounting for the known 80% deuterium content of the deuterium exchange buffer.

Peptides were identified using tandem MS (MS/MS) with a Fusion Orbitrap mass spectrometer (ThermoFisher). Product ion spectra were acquired in data-dependent mode with the top eight most abundant ions selected for the product ion analysis per scan event. The MS/MS data files were submitted to Proteome Discover 2.4 (ThermoFisher) for high confident peptide identification.

The HDX data from all overlapping peptides were consolidated to individual amino acid values using a residue averaging approach. Briefly, for each residue, the deuterium incorporation values and peptide lengths from all overlapping peptides were assembled. A weighting function was applied in which shorter peptides were weighted more heavily and longer peptides were weighted less. Each of the weighted deuterium incorporation values were then averaged to produce a single value for each amino acid. The initial two residues of each peptide, as well as prolines, were omitted from the calculations.

**Fig. S1. Purification and characterization of the Omicron spike protein complex.** (**A**) Gel filtration profile of the Omicron ECD-ACE2 complex, showing a sharp peak, and the SDS gel of the Omicron ECD-ACE2 complex, showing balanced ratios for each subunit. (**B**) Gel filtration profile of the Omicron ECD-Fab complex, showing a sharp peak, and the SDS gel of the Omicron ECD-Fab complex, showing balanced ratios for each subunit. (**C**) Thermal stability shift analysis of the Omicron and WT spike trimer. (**D**) Thermal stability shift analysis of the Omicron and WT RBD.

**Fig. S2. Cryo-EM data processing of the Omicron ECD-ACE2 complex.** (**A**) A representative cryo-EM micrograph of Omicron ECD-ACE2 complex with 50 nm scale bar included as a size reference. (**B**) Computational processing of cryo-EM data. (**C**) Twenty representative reference-free two-dimensional (2D) cryo-EM class averages reveal the ACE2 density (yellow arrow). (**D**) Local resolution of sub-reconstructions of RBD-ACE2 (left panel) and RBD-ACE2-RBD (right panel). Resolution bar is shown in the middle. (**E**) The FSC curves for the reconstructions. Color scheme: ECD-ACE2, black; RBD-ACE2, red; ECD-trimer, yellow; RBD-ACE2-RBD, blue. The resolution of the reconstructions using the Fourier shell cutoff at 0.143 is shown.

**Fig. S3. Cryo-EM analysis of Omicron ECD-trimer.** (**A**) High resolution density map of Omicron ECD-trimer after sharpening. Front and top views are shown. The densities for three RBDs are smeared. (**B**) Density map of Omicron ECD-trimer at low resolution without sharpening. The blob densities according to RBDs are shown in all-down conformation. Front and top views are shown. Three ECDs are colored in purple, green, and salmon.

**Fig. S4** Comparative HDX analysis on WT vs Omicron Spike protein. (**A**) Single amide consolidated HDX-MS profiling of *apo* WT Spike vs apo Omicron full-length Spike protein. The y- and x-axes illustrate the differential D% ($HDX_{Omicron}$ minus

$HDX_{WT}$) and residue numbers of aligned WT and Omicron Spike protein, respectively. (**B**) Differential HDX data consolidated from Fig. S1 and Fig.S2 are mapped to Full-length Omicron spike conformer that displays an up-oriented RBD, according to the differential HDX dynamics key shown below. Regions that show increased HDX activity (more disordered) are colored orange/red; regions that show decreased HDX activity (more stable) are colored blue.

**Fig. S5 Neutralization of WT, Alpha, Beta, Gamma, and Delta pseudo-virus by JMB2002.** Delta IC50 values were calculated by Prism V8.0 software using a four-parameter logistic curve fitting approach.

**Fig. S6. Cryo-EM data processing of the Omicron ECD-Fab complex.** (**A**) Flow chart of cryo-EM analysis. (**B**) Representative cryo-EM image (scale bar, 50 nm) from 14,740 movies. (**C**) Representative 2D averages (scale bar, 5 nm). (**D**) The focus refinement density map of the binding interfaces between RBDs and Fabs. (**E**) Local resolution for the densities. (**F-G**) ‘Gold-standard’ FSC curves of the global complex and the RBDs-Fabs interface.

**Fig. S7. Structural comparisons of the ACE2-bound and Fab-bound SARS-CoV-2 Omicron Spike protein.** (**A-B**) Superpositions of the ACE2-bound and Fab-bound Omicron S trimer. The different protomers are colored. The compared models showed with a front view (A) and a top view (B). (**C**) Superpositions shows ACE2 and the Fab do not share binding sites of S protein RBD.

**Fig. S8. Different states of the Omicron ECD-Fab complexes.** (**A**) The cryo-EM density map and model of 3 Fab-bound ECD-Fab complex showed in front view and top view. (**B**) The cryo-EM density map and model of 1 Fab-bound ECD-Fab complex showed in front view and top view.

### Fig. 1

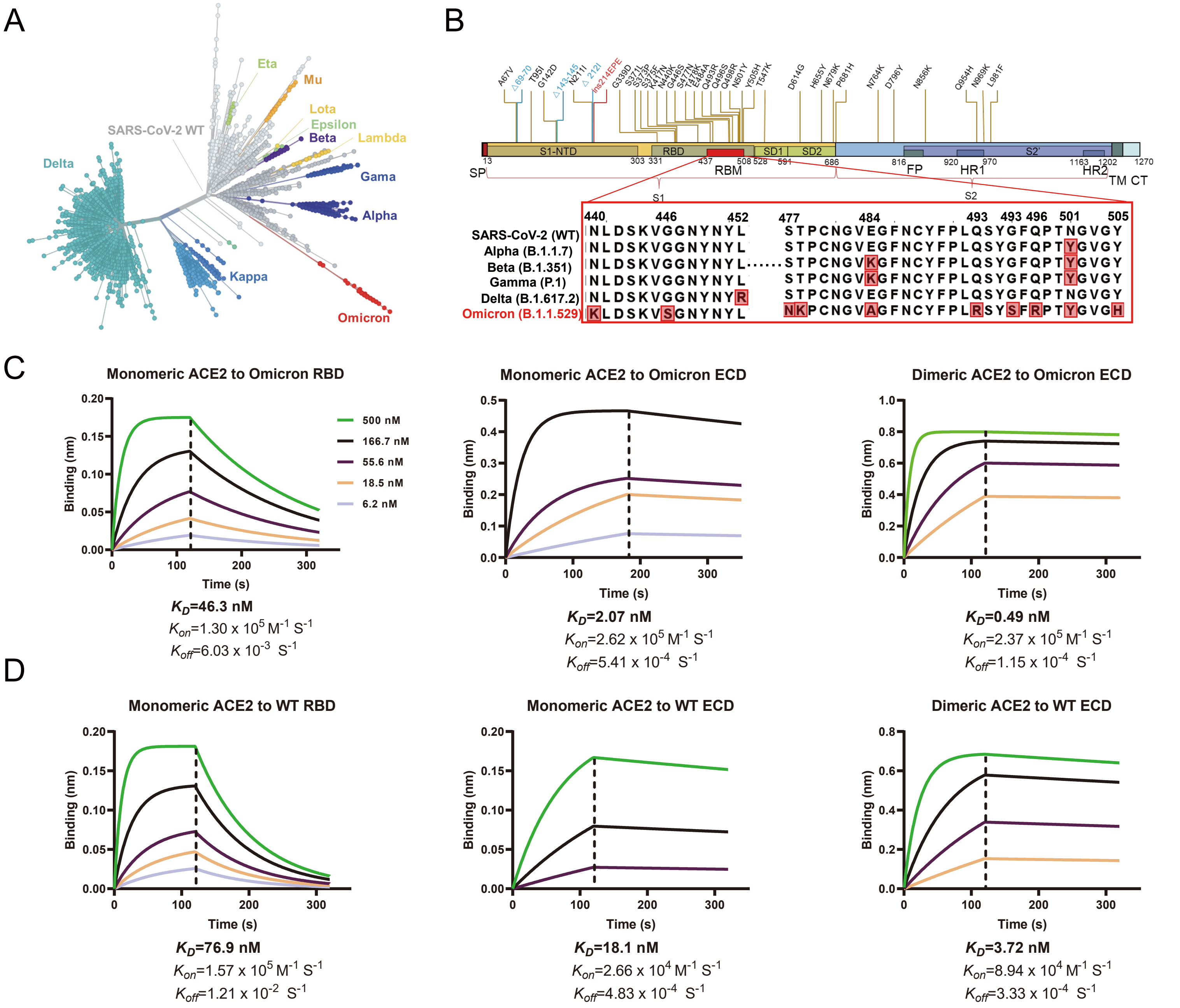
A
Eta
Mu
Lota
Epsilon
Beta
Lambda
SARS-CoV-2 WT
Delta
Gama
Alpha
Kappa
Omicron
B
A67V
Δ69-70
T95I
G142D
Δ143-145
N211I
Δ212I
ins214EPE
G339D
S371L
S373P
S375F
K417N
N440K
G446S
S477N
T478K
E484A
Q493R
G496S
Q498R
N501Y
Y505H
T547K
D614G
H655Y
N679K
P681H
N764K
D796Y
N856K
Q954H
N969K
L981F
S1-NTD
RBD
SD1
SD2
S2'
13 303 331 437 508 528 591 686 816 920 970 1163 1202 1270
SP
RBM
FP
HR1
HR2
TM CT
S1
S2
440 446 452 477 484 493 493 496 501 505
SARS-CoV-2 (WT) NLDSKVGGNYNYL STPCNGVEGFNCYFPLQSYGFQPTNGVGY
Alpha (B.1.1.7) NLDSKVGGNYNYL STPCNGVEGFNCYFPLQSYGFQPTYGVGY
Beta (B.1.351) NLDSKVGGNYNYL STPCNGVKGFNCYFPLQSYGFQPTYGVGY
Gamma (P.1) NLDSKVGGNYNYL STPCNGVKGFNCYFPLQSYGFQPTYGVGY
Delta (B.1.617.2) NLDSKVGGNYNYR STPCNGVEGFNCYFPLQSYGFQPTNGVGY
Omicron (B.1.1.529) KLDSKVSGNYNYL NKPCNGVAGFNCYFPLRSYSFRPTYGVGH
C
Monomeric ACE2 to Omicron RBD
500 nM
166.7 nM
55.6 nM
18.5 nM
6.2 nM
Binding (nm)
Time (s)
$K_D$=46.3 nM
$K_{on}$=1.30 x $10^5$ $M^{-1}$ $S^{-1}$
$K_{off}$=6.03 x $10^{-3}$ $S^{-1}$
Monomeric ACE2 to Omicron ECD
$K_D$=2.07 nM
$K_{on}$=2.62 x $10^5$ $M^{-1}$ $S^{-1}$
$K_{off}$=5.41 x $10^{-4}$ $S^{-1}$
Dimeric ACE2 to Omicron ECD
$K_D$=0.49 nM
$K_{on}$=2.37 x $10^5$ $M^{-1}$ $S^{-1}$
$K_{off}$=1.15 x $10^{-4}$ $S^{-1}$
D
Monomeric ACE2 to WT RBD
$K_D$=76.9 nM
$K_{on}$=1.57 x $10^5$ $M^{-1}$ $S^{-1}$
$K_{off}$=1.21 x $10^{-2}$ $S^{-1}$
Monomeric ACE2 to WT ECD
$K_D$=18.1 nM
$K_{on}$=2.66 x $10^4$ $M^{-1}$ $S^{-1}$
$K_{off}$=4.83 x $10^{-4}$ $S^{-1}$
Dimeric ACE2 to WT ECD
$K_D$=3.72 nM
$K_{on}$=8.94 x $10^4$ $M^{-1}$ $S^{-1}$
$K_{off}$=3.33 x $10^{-4}$ $S^{-1}$

Fig. 2

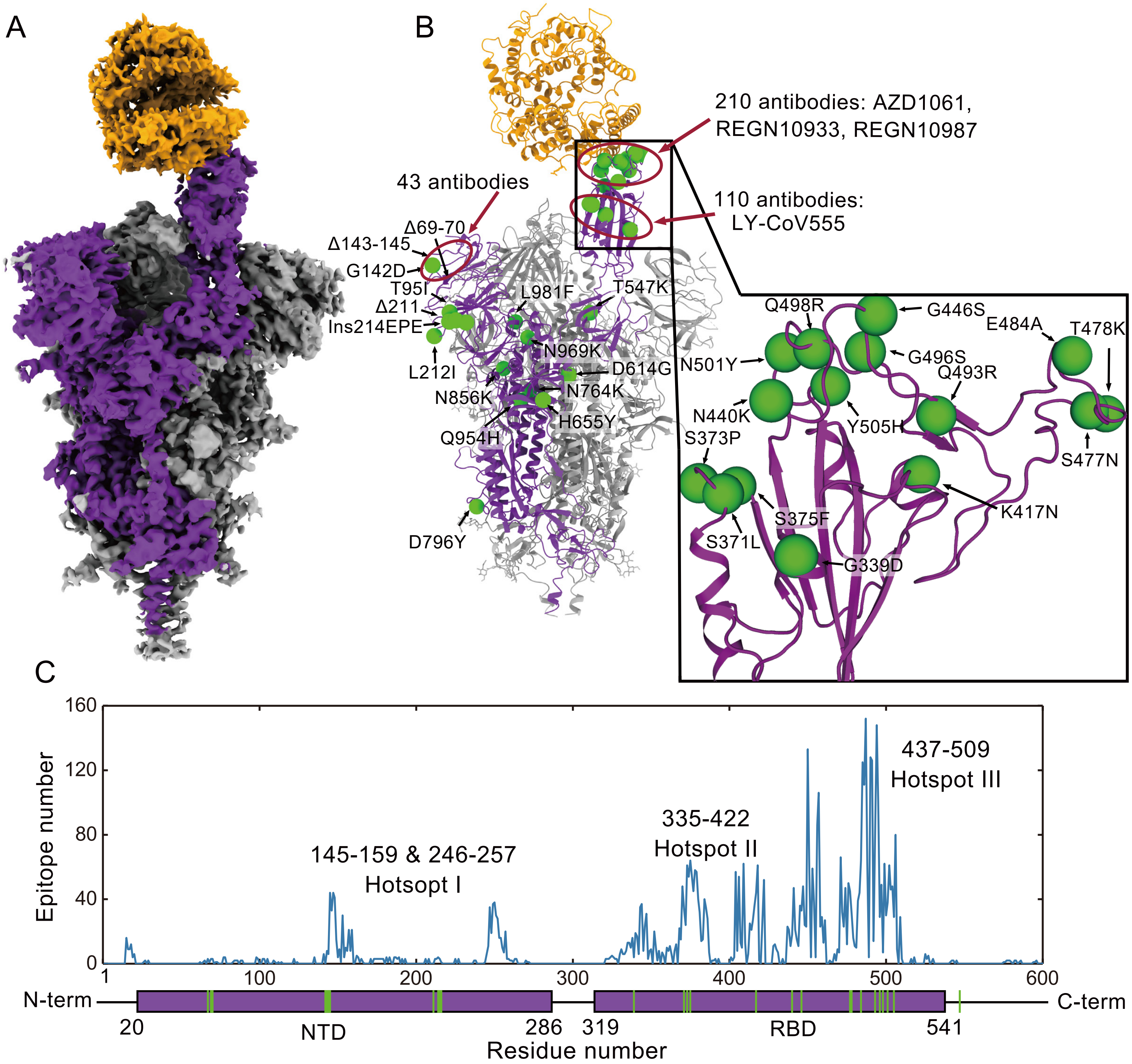
A
B
210 antibodies: AZD1061, REGN10933, REGN10987
43 antibodies
110 antibodies: LY-CoV555
Δ69-70
Δ143-145
G142D
T95I
Δ211
Ins214EPE
L212I
L981F
T547K
N969K
D614G
N764K
N856K
H655Y
Q954H
D796Y
Q498R
G446S
E484A
T478K
N501Y
G496S
Q493R
N440K
Y505H
S373P
S477N
S375F
K417N
S371L
G339D
C
Epitope number
145-159 & 246-257
Hotsopt I
335-422
Hotspot II
437-509
Hotspot III
N-term
C-term
NTD
RBD
20
286
319
541
Residue number

Fig. 3

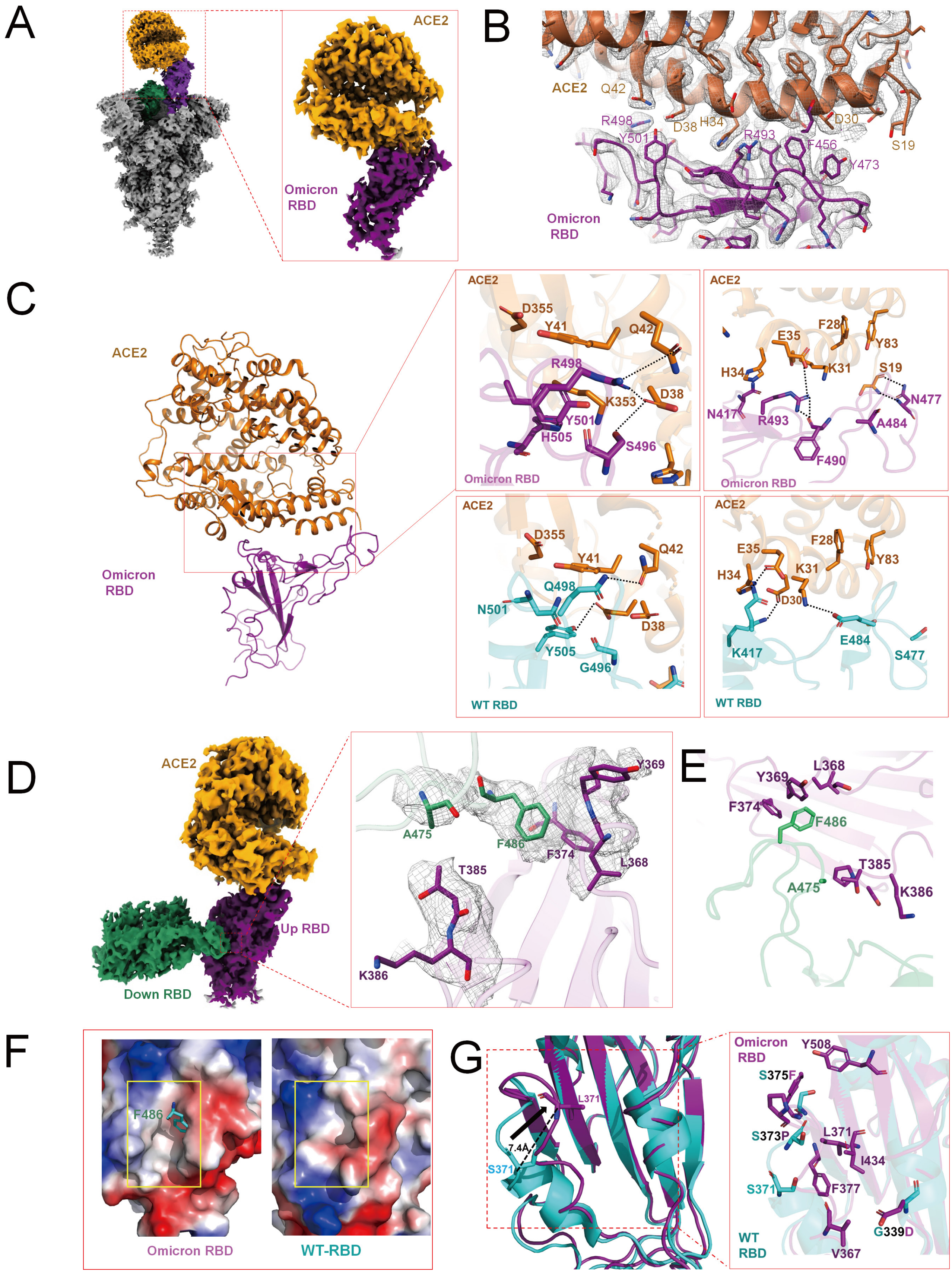
A
ACE2
Omicron RBD
B
ACE2
Q42
R498
D38
H34
Y501
R493
F456
D30
S19
Y473
Omicron RBD
C
ACE2
Omicron RBD
D355
Y41
Q42
R498
K353
D38
Y501
H505
S496
F28
E35
Y83
K31
H34
S19
N477
N417
R493
A484
F490
WT RBD
D355
Y41
Q42
Q498
N501
D38
Y505
G496
F28
E35
Y83
K31
H34
D30
K417
E484
S477
D
ACE2
Up RBD
Down RBD
Y369
A475
F486
F374
L368
T385
K386
E
Y369
L368
F374
F486
T385
A475
K386
F
F486
Omicron RBD
WT-RBD
G
L371
7.4Å
S371
Omicron RBD
Y508
S375F
S373P
L371
I434
S371
F377
WT RBD
G339D
V367

Fig. 4

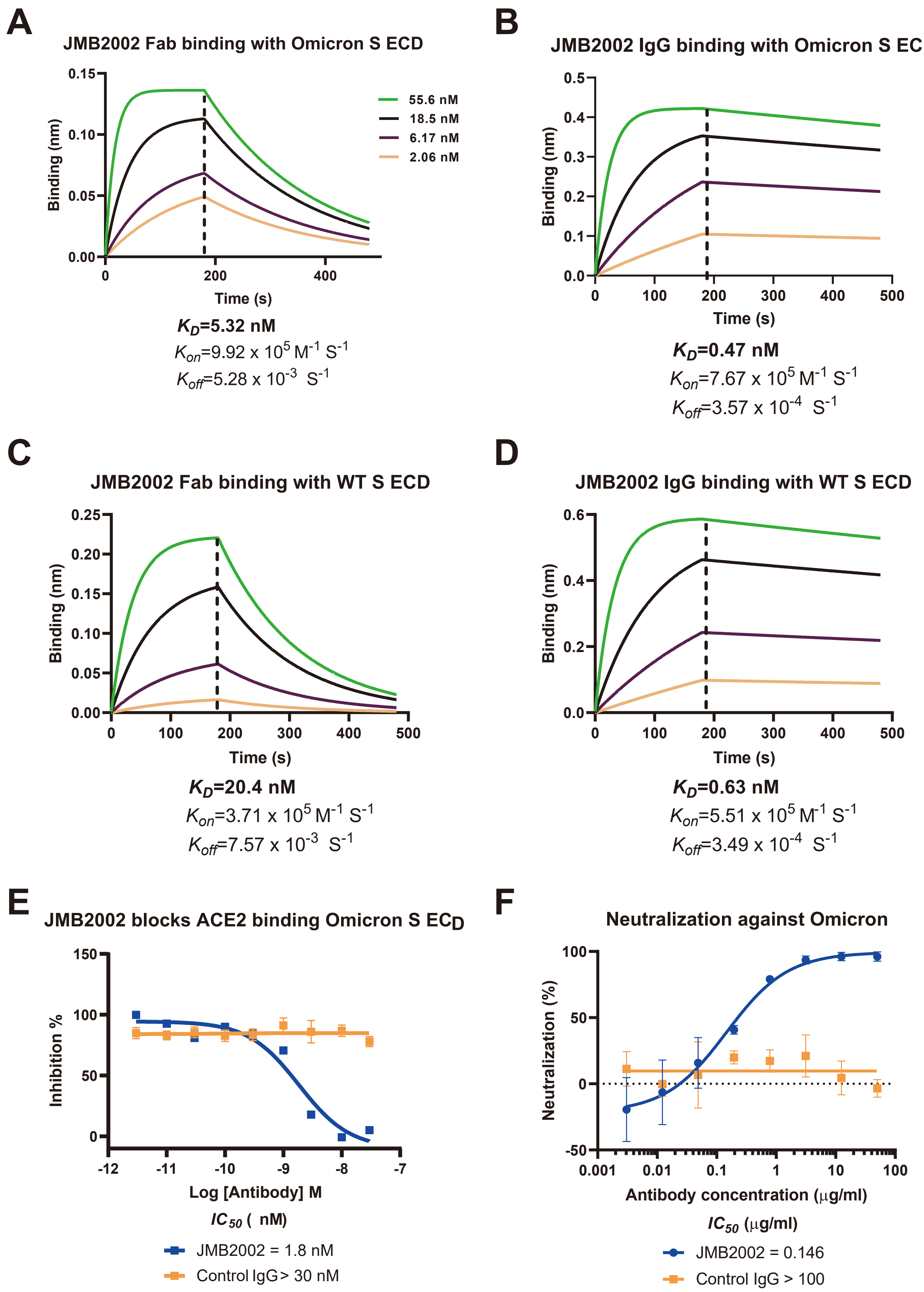
A
JMB2002 Fab binding with Omicron S ECD
Binding (nm)
Time (s)
55.6 nM
18.5 nM
6.17 nM
2.06 nM
$K_D$=5.32 nM
$K_{on}$=9.92 x $10^5$ $M^{-1}$ $S^{-1}$
$K_{off}$=5.28 x $10^{-3}$ $S^{-1}$
B
JMB2002 IgG binding with Omicron S ECD
Binding (nm)
Time (s)
$K_D$=0.47 nM
$K_{on}$=7.67 x $10^5$ $M^{-1}$ $S^{-1}$
$K_{off}$=3.57 x $10^{-4}$ $S^{-1}$
C
JMB2002 Fab binding with WT S ECD
Binding (nm)
Time (s)
$K_D$=20.4 nM
$K_{on}$=3.71 x $10^5$ $M^{-1}$ $S^{-1}$
$K_{off}$=7.57 x $10^{-3}$ $S^{-1}$
D
JMB2002 IgG binding with WT S ECD
Binding (nm)
Time (s)
$K_D$=0.63 nM
$K_{on}$=5.51 x $10^5$ $M^{-1}$ $S^{-1}$
$K_{off}$=3.49 x $10^{-4}$ $S^{-1}$
E
JMB2002 blocks ACE2 binding Omicron S ECD
Inhibition %
Log [Antibody] M
$IC_{50}$ ( nM)
JMB2002 = 1.8 nM
Control IgG> 30 nM
F
Neutralization against Omicron
Neutralization (%)
Antibody concentration (μg/ml)
$IC_{50}$ (μg/ml)
JMB2002 = 0.146
Control IgG > 100

Fig. 5

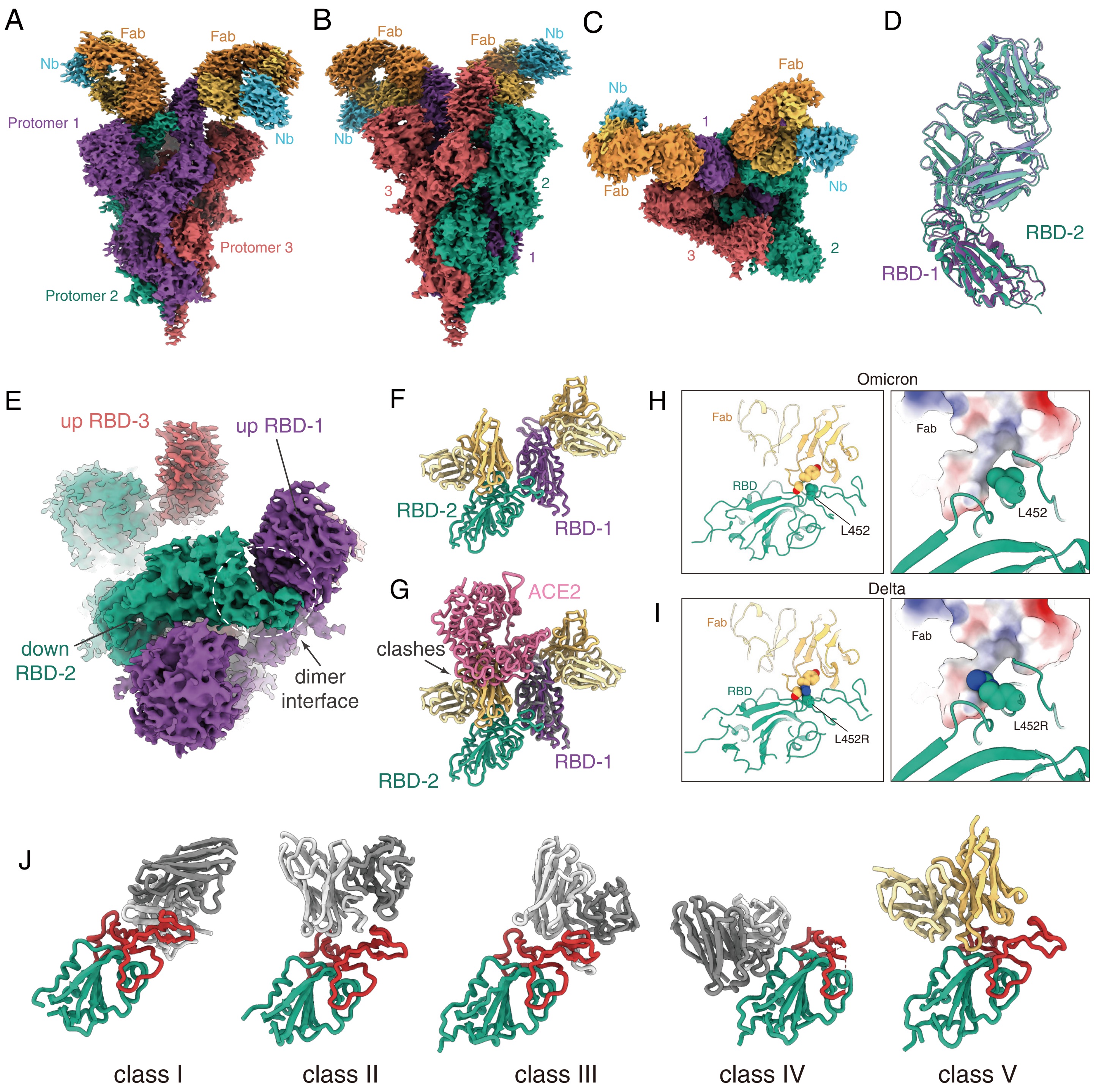

A
Fab
Fab
Nb
Protomer 1
Nb
Protomer 3
Protomer 2
B
Fab
Fab
Nb
Nb
3
2
1
C
Fab
Nb
1
Fab
Nb
3
2
D
RBD-2
RBD-1
E
up RBD-3
up RBD-1
down RBD-2
dimer interface
F
RBD-2
RBD-1
G
ACE2
clashes
RBD-1
RBD-2
H
Omicron
Fab
RBD
L452
Fab
L452
I
Delta
Fab
RBD
L452R
Fab
L452R
J
class I
class II
class III
class IV
class V

### Fig. S1

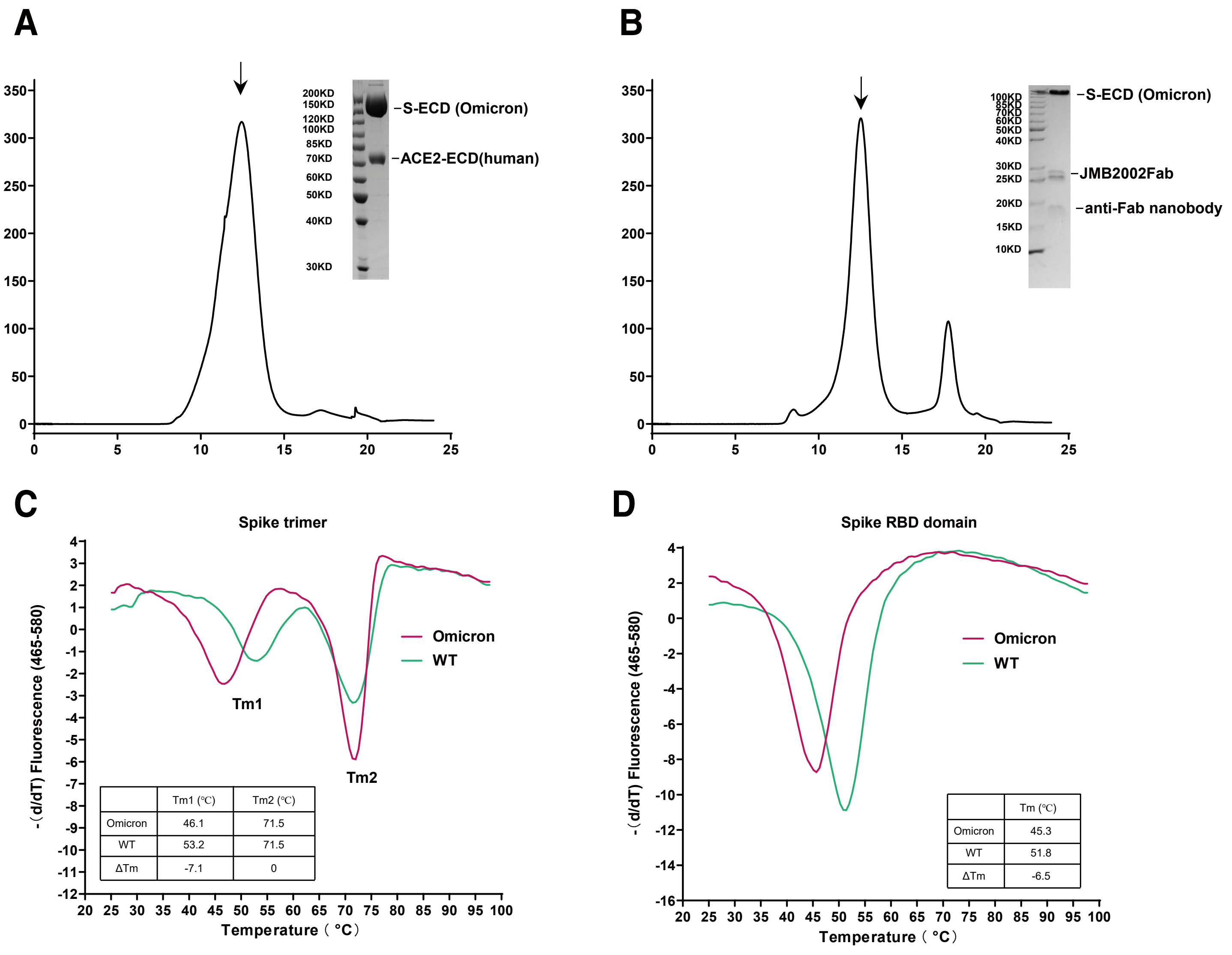
A
200KD
150KD
120KD
100KD
85KD
70KD
60KD
50KD
40KD
30KD
S-ECD (Omicron)
ACE2-ECD(human)
B
100KD
85KD
70KD
60KD
50KD
40KD
30KD
25KD
20KD
15KD
10KD
S-ECD (Omicron)
JMB2002Fab
anti-Fab nanobody
C
Spike trimer
Omicron
WT
Tm1
Tm2
-（d/dT) Fluorescence (465-580)
Temperature（°C）
Tm1 (℃) | Tm2 (℃)
Omicron | 46.1 | 71.5
WT | 53.2 | 71.5
ΔTm | -7.1 | 0
D
Spike RBD domain
Omicron
WT
-（d/dT) Fluorescence (465-580)
Temperature（°C）
Tm (℃)
Omicron | 45.3
WT | 51.8
ΔTm | -6.5

Fig. S2

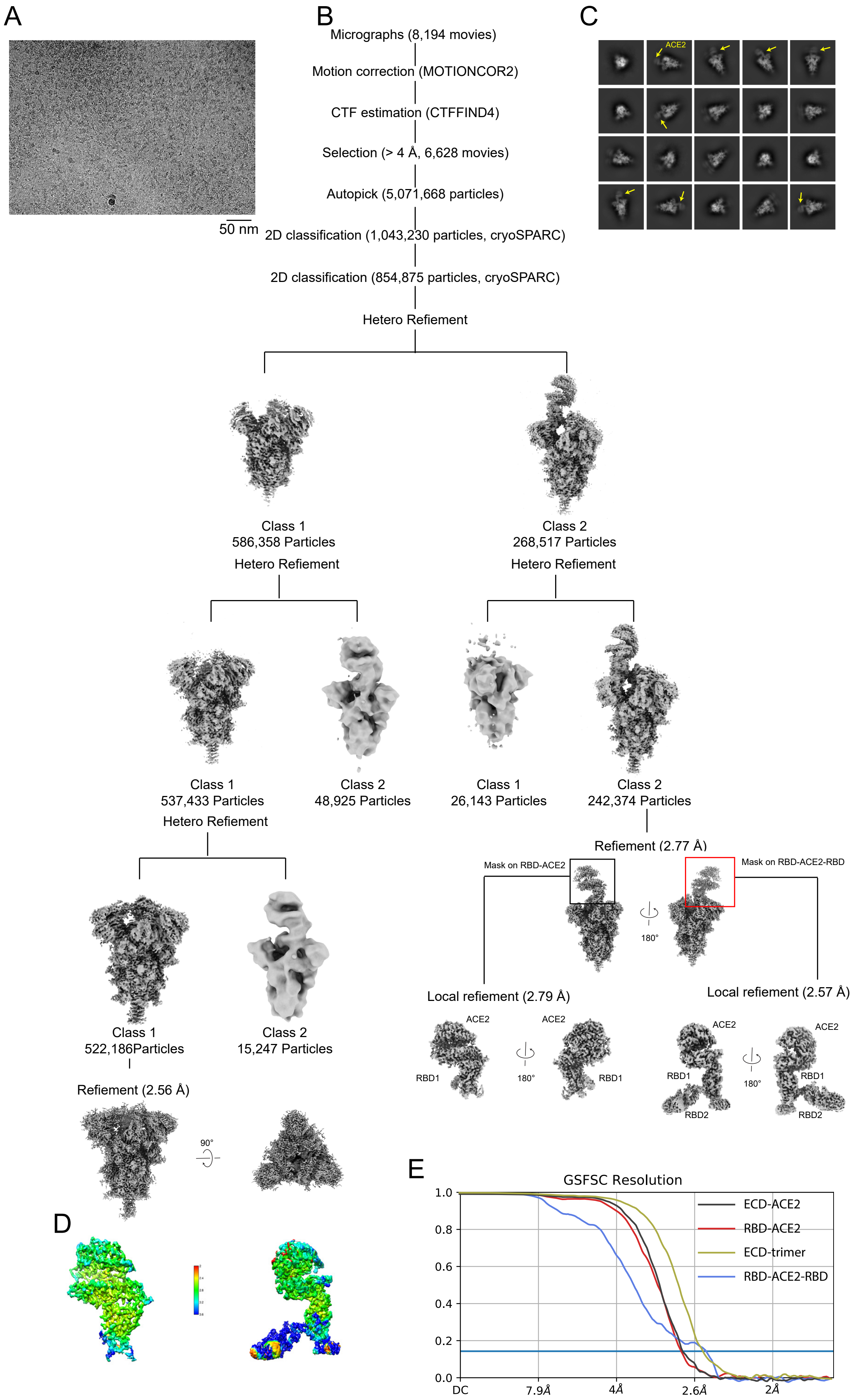
A
50 nm
B
Micrographs (8,194 movies)
Motion correction (MOTIONCOR2)
CTF estimation (CTFFIND4)
Selection (> 4 Å, 6,628 movies)
Autopick (5,071,668 particles)
2D classification (1,043,230 particles, cryoSPARC)
2D classification (854,875 particles, cryoSPARC)
Hetero Refiement
Class 1
586,358 Particles
Hetero Refiement
Class 2
268,517 Particles
Hetero Refiement
Class 1
537,433 Particles
Hetero Refiement
Class 2
48,925 Particles
Class 1
26,143 Particles
Class 2
242,374 Particles
Refiement (2.77 Å)
Mask on RBD-ACE2
Mask on RBD-ACE2-RBD
180°
Class 1
522,186Particles
Class 2
15,247 Particles
Local refiement (2.79 Å)
Local refiement (2.57 Å)
ACE2
RBD1
RBD2
Refiement (2.56 Å)
90°
C
ACE2
D
E
GSFSC Resolution
ECD-ACE2
RBD-ACE2
ECD-trimer
RBD-ACE2-RBD
1.0
0.8
0.6
0.4
0.2
0.0
DC
7.9Å
4Å
2.6Å
2Å

### Fig. S3

## A

Sharpened high resolution map (2.56 Å, Bfactor=92)

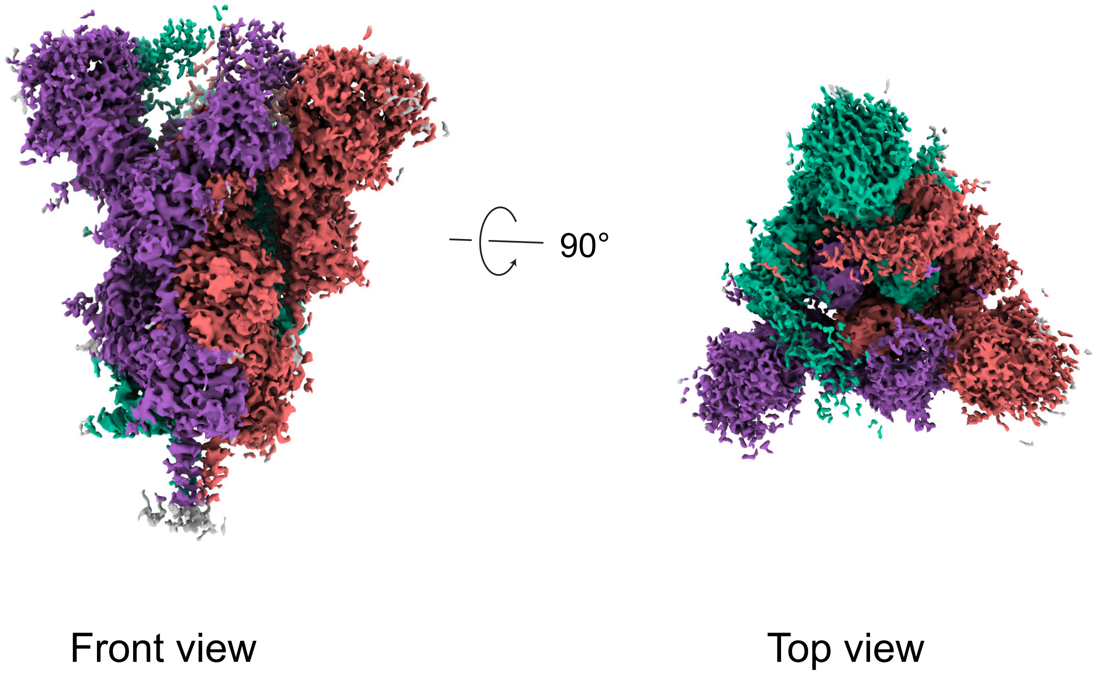


## B

Unsharpened low resolution map (6.5 Å)

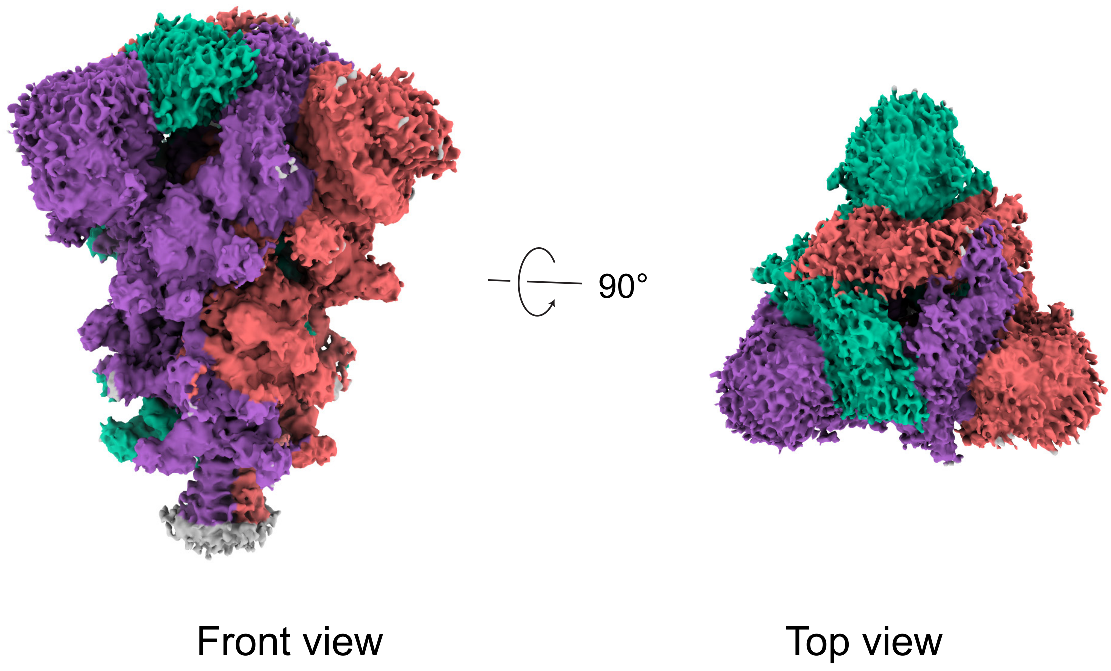

### Fig. S4

A

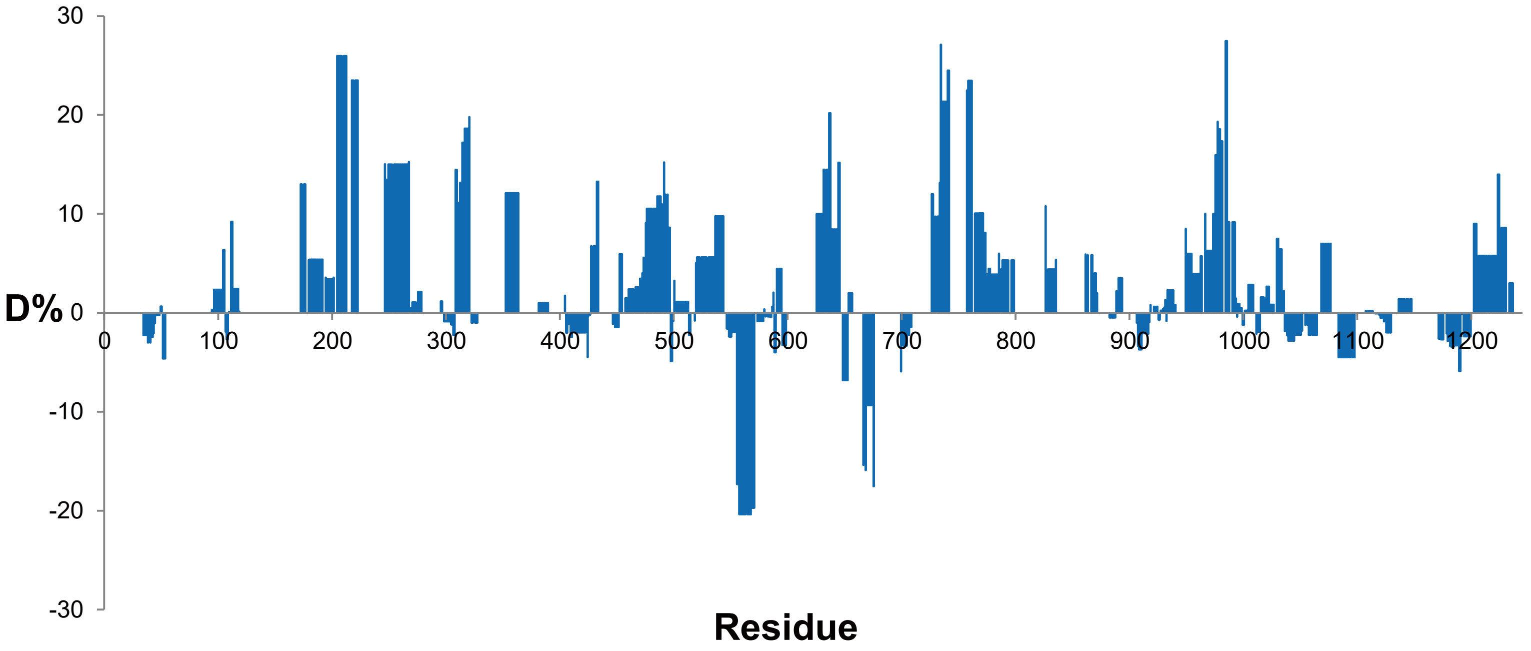
30
20
10
D% 0
-10
-20
-30
0
100
200
300
400
500
600
700
800
900
1000
1100
1200
Residue

B

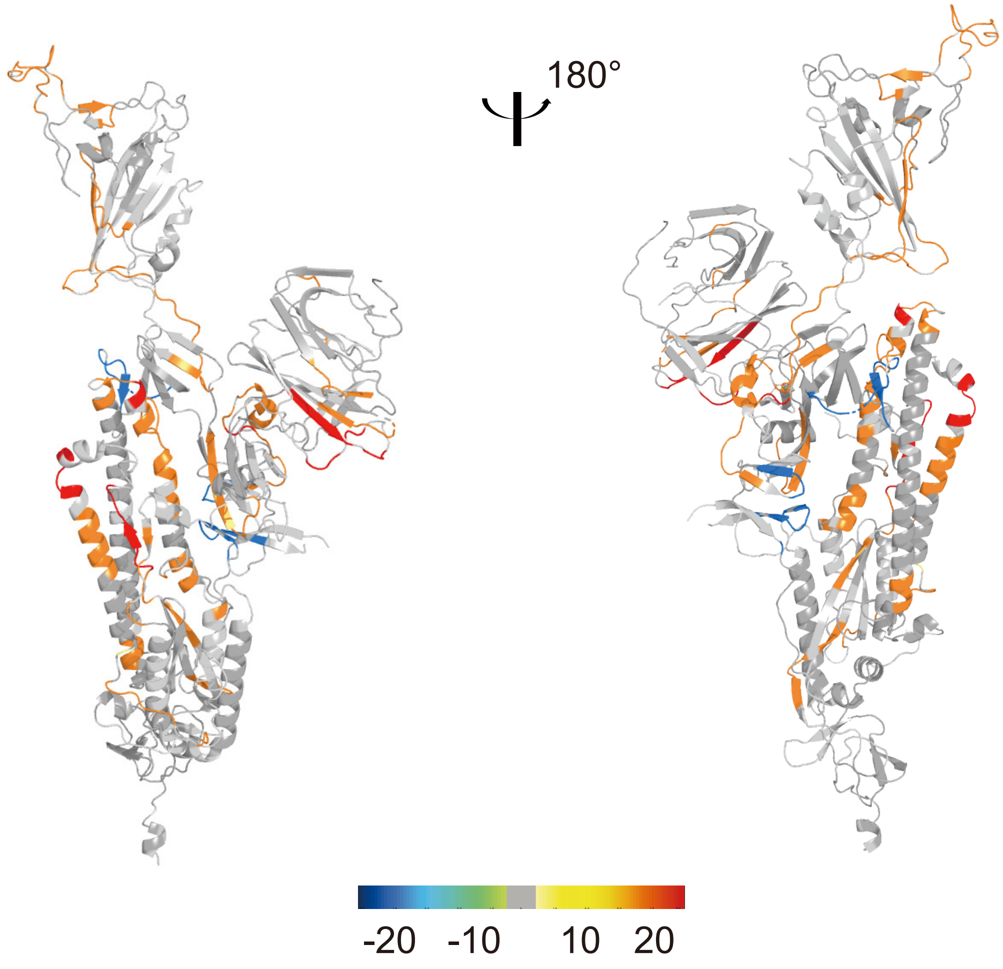
180°
-20
-10
10
20

Fig. S5

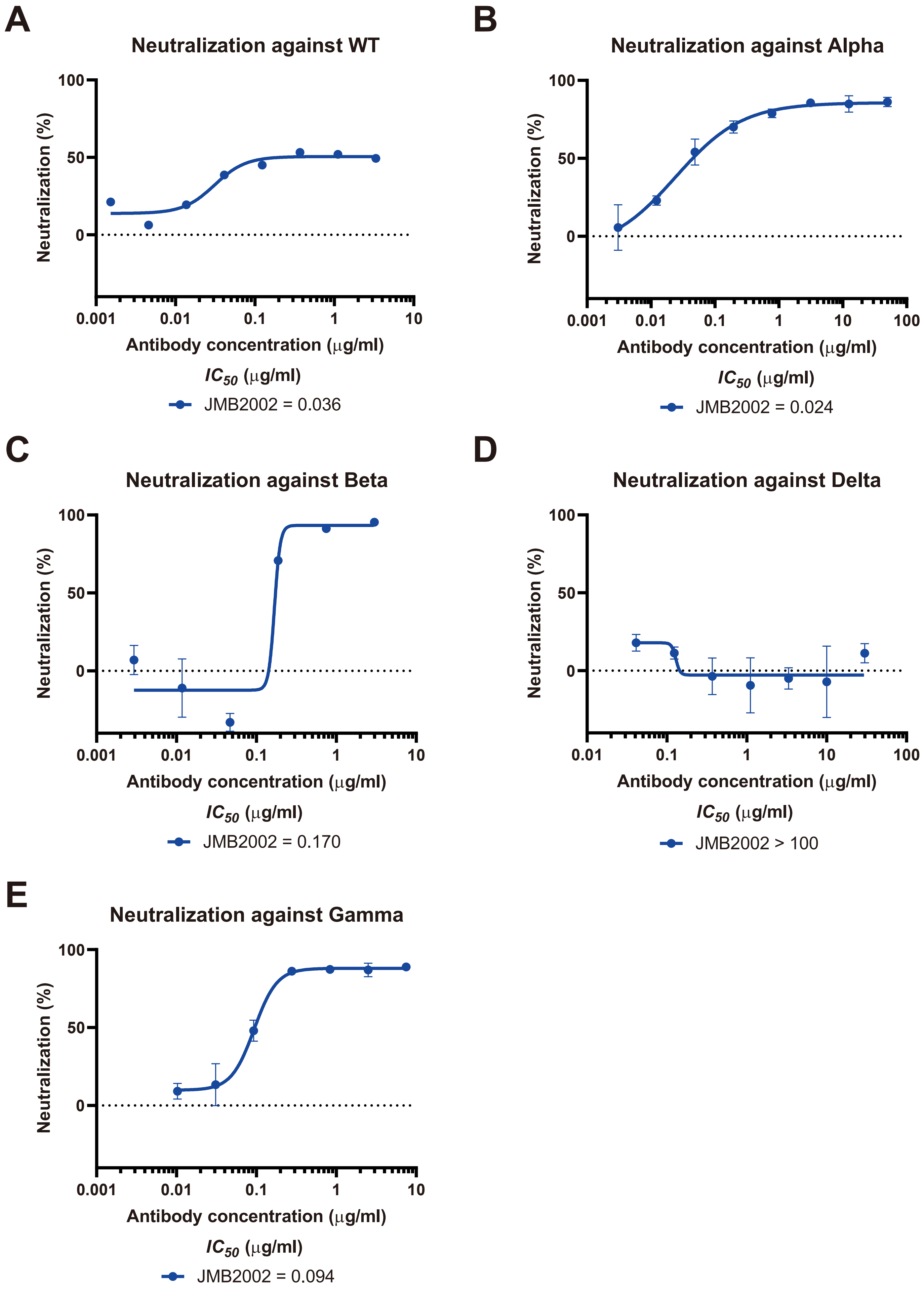
A
Neutralization against WT
Neutralization (%)
100
50
0
0.001
0.01
0.1
1
10
Antibody concentration (μg/ml)
IC50 (μg/ml)
JMB2002 = 0.036
B
Neutralization against Alpha
Neutralization (%)
100
50
0
0.001
0.01
0.1
1
10
100
Antibody concentration (μg/ml)
IC50 (μg/ml)
JMB2002 = 0.024
C
Neutralization against Beta
Neutralization (%)
100
50
0
0.001
0.01
0.1
1
10
Antibody concentration (μg/ml)
IC50 (μg/ml)
JMB2002 = 0.170
D
Neutralization against Delta
Neutralization (%)
100
50
0
0.01
0.1
1
10
100
Antibody concentration (μg/ml)
IC50 (μg/ml)
JMB2002 > 100
E
Neutralization against Gamma
Neutralization (%)
100
50
0
0.001
0.01
0.1
1
10
Antibody concentration (μg/ml)
IC50 (μg/ml)
JMB2002 = 0.094

Fig. S6

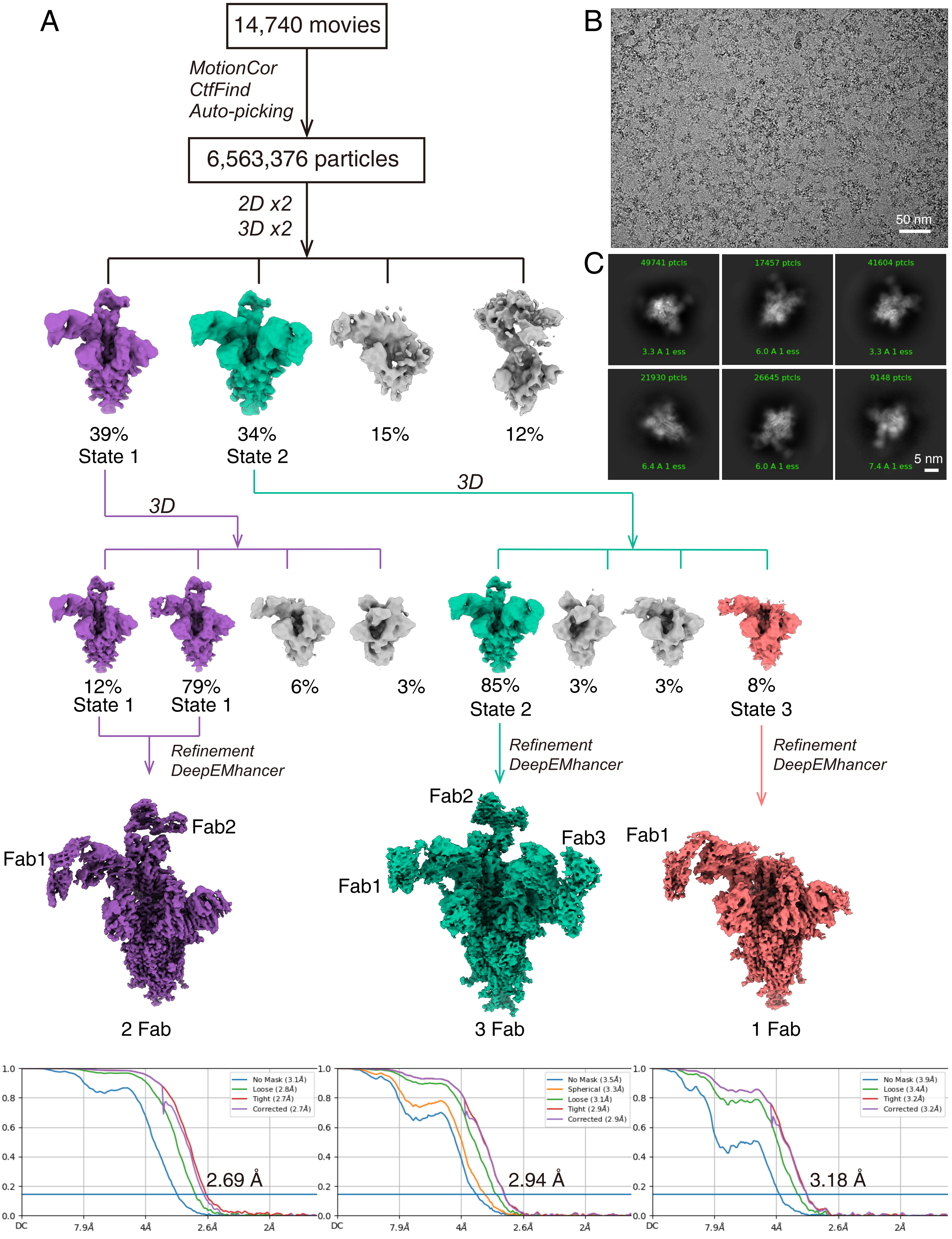

A
14,740 movies
MotionCor
CtfFind
Auto-picking
6,563,376 particles
2D x2
3D x2
39%
State 1
34%
State 2
15%
12%
3D
3D
12%
State 1
79%
State 1
6%
3%
85%
State 2
3%
3%
8%
State 3
Refinement
DeepEMhancer
Refinement
DeepEMhancer
Refinement
DeepEMhancer
Fab1
Fab2
2 Fab
Fab2
Fab3
Fab1
3 Fab
Fab1
1 Fab
B
50 nm
C
49741 ptcls
17457 ptcls
41604 ptcls
3.3 A 1 ess
6.0 A 1 ess
3.3 A 1 ess
21930 ptcls
26645 ptcls
9148 ptcls
6.4 A 1 ess
6.0 A 1 ess
7.4 A 1 ess
5 nm
No Mask (3.1Å)
Loose (2.8Å)
Tight (2.7Å)
Corrected (2.7Å)
2.69 Å
No Mask (3.5Å)
Spherical (3.3Å)
Loose (3.1Å)
Tight (2.9Å)
Corrected (2.9Å)
2.94 Å
No Mask (3.9Å)
Loose (3.4Å)
Tight (3.2Å)
Corrected (3.2Å)
3.18 Å
DC
7.9Å
4Å
2.6Å
2Å

Fig. S7

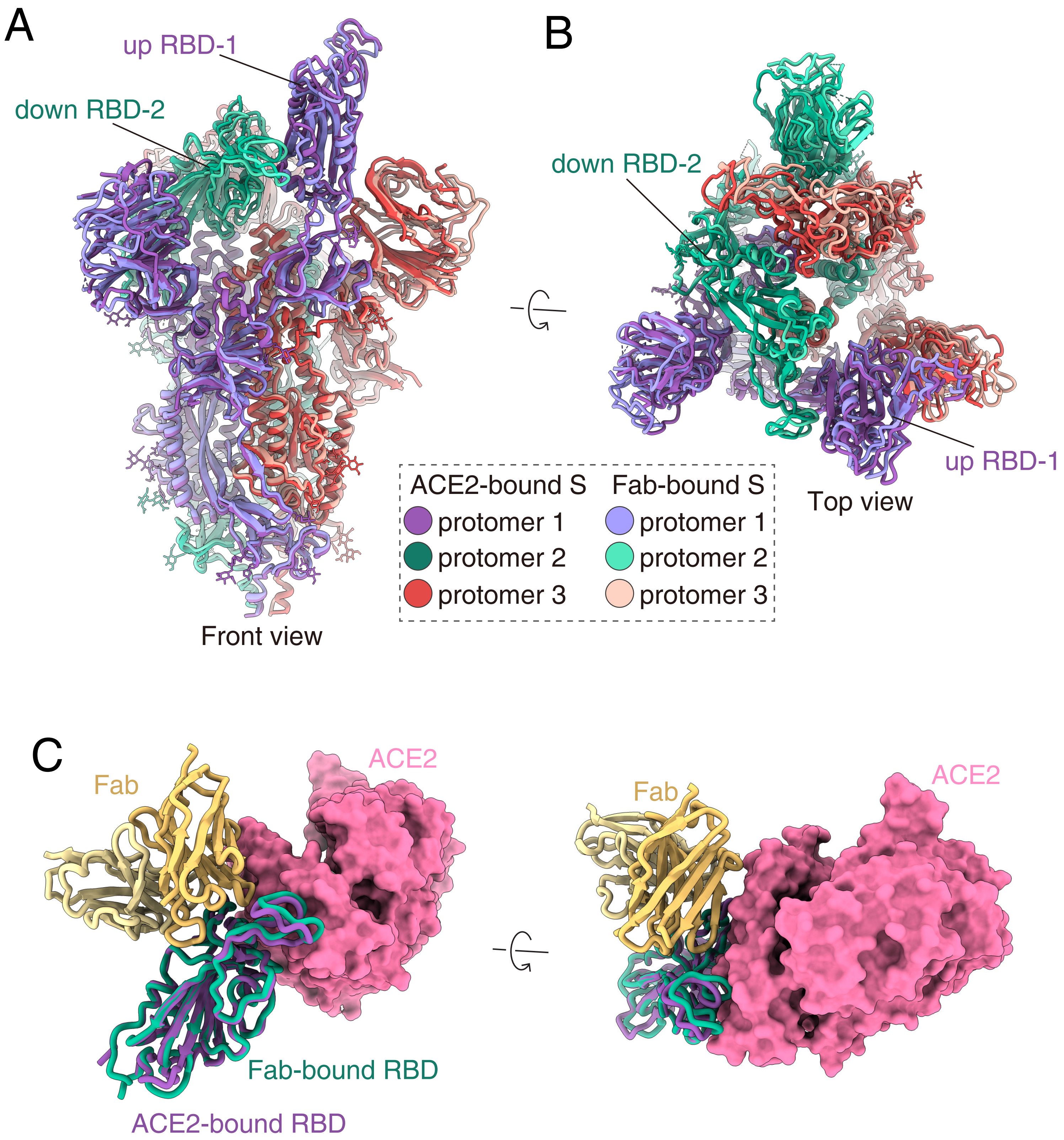
A
up RBD-1
down RBD-2
Front view
B
down RBD-2
up RBD-1
Top view
ACE2-bound S
protomer 1
protomer 2
protomer 3
Fab-bound S
protomer 1
protomer 2
protomer 3
C
Fab
ACE2
Fab-bound RBD
ACE2-bound RBD
Fab
ACE2

### Fig. S8

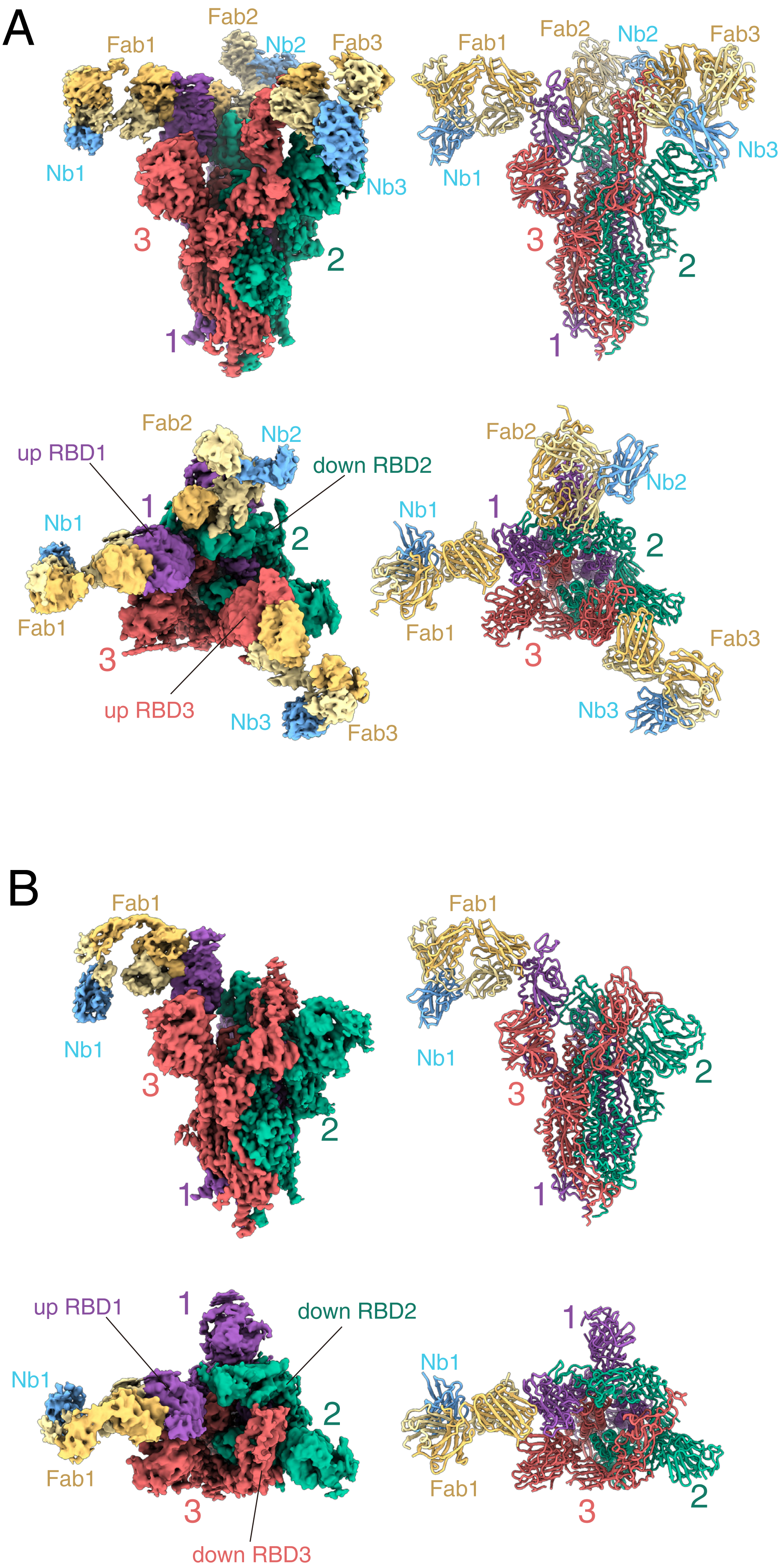
A
Fab1
Fab2
Nb2
Fab3
Nb1
Nb3
3
2
1
up RBD1
down RBD2
up RBD3
B
Fab1
Nb1
3
2
1
up RBD1
down RBD2
down RBD3

### Table S1 Cryo-EM data collection, refinement and validation statistics

| | Omicron ECD-ACE2 complex | | | | Omicron ECD-Fab complex | | |
|---|---|---|---|---|---|---|---|
| **Data collection and processing** | | | | | | | |
| Magnification | 105,000 | | | | 105,000 | | |
| Voltage (kV) | 300 | | | | 300 | | |
| Electron exposure ($e^-/Å^2$) | 50 | | | | 50 | | |
| Defocus range (μm) | -0.5 ~ -2.5 | | | | -0.5 ~ -2.5 | | |
| Pixel size (Å) | 0.824 | | | | 0.824 | | |
| Symmetry imposed | C1 | | | | C1 | | |
| Initial particle projections (no.) | 5,071,668 | | | | 6,563,376 | | |
| Sub-maps | Map 1 ECD-Trimer | Map 2 ECD-ACE2 | Map 3 RBD-ACE2 | Map 4 RBD-ACE2-RBD | Map 1 ECD-2Fab | Map 2 RBD-3Fab | Map 3 ECD-1Fab |
| Final particle projections (no.) | 522,186 | 242,374 | 242,374 | 242,374 | 906,588 | 434,893 | 172,001 |
| Map resolution (Å) | 2.56 | 2.77 | 2.79 | 2.56 | 2.69 | 2.92 | 3.18 |
| FSC threshold | 0.143 | 0.143 | 0.143 | 0.143 | 0.143 | 0.143 | 0.143 |
| Map resolution range (Å) | 2.6-5.5 | 2.8-6.2 | 2.8-3.2 | 2.6-5.5 | 2.5-6.0 | 2.5-6.0 | 2.5-6.0 |
| **Refinement** | | | | | | | |
| Initial model used | 6VXX | 7A94 | 7A94 | 7A94 | 6VXX | 6VXX | 6VXX |
| Model resolution (Å) | 3.1 | 3.9 | 3.9 | 3.9 | 2.8 | 3.0 | 3.2 |
| FSC threshold | 0.143 | 0.143 | 0.143 | 0.143 | 0.143 | 0.143 | 0.143 |
| Map sharpening *B* factor ($Å^2$) | -92 | -91.3 | -107.1 | -64.6 | DeepEMhancer | | |
| Model composition | | | | | | | |
| Non-hydrogen atoms | | | 6491 | 8081 | 28,068 | 32,295 | 36,414 |
| Protein residues | | | 793 | 996 | 3657 | 4216 | 4785 |
| R.m.s. deviations | | | | | | | |
| Bond lengths (Å) | | | 0.005 | 0.007 | 0.013 | 0.013 | 0.015 |
| Bond angles (°) | | | 0.873 | 1.10 | 1.389 | 1.290 | 1.539 |
| Validation | | | | | | | |
| MolProbity score | | | 1.39 | 0 | 1.08 | 1.05 | 1.13 |
| Clashscore | | | 3.63 | 3.67 | 0.07 | 0.03 | 0.10 |
| Rotamer outliers (%) | | | 0.00 | 0.34 | 0.45 | 0.50 | 0.10 |
| Ramachandran plot | | | | | | | |
| Favored (%) | | | 96.45 | 94.42 | 93.90 | 94.04 | 93.08 |
| Allowed (%) | | | 3.55 | 5.58 | 6.10 | 5.96 | 6.92 |
| Disallowed (%) | | | 0 | 0 | 0 | 0 | 0 |

**Table S2**

**A summary of the comparison between our ACE2-RBD$^{Omi}$ structure and selected ACE2-RBD structures.**

| Our structure Compared with | | 6M0J | 6LZG | 6M17 | 7A94 | 7KMB | 7DF4 |
|---|---|---|---|---|---|---|---|
| Method | | X-ray | X-ray | Cryo-EM | Cryo-EM | Cryo-EM | Cryo-EM |
| Resolution (Å) | | 2.45 | 2.5 | 2.9 | 3.4 | 3.39 | 3.8 |
| RMSD (Å) | Overall | 0.49 | 0.44 | 0.60 | 0.87 | 1.18 | 1.34 |
| | ACE2 | 0.49 | 0.44 | 0.60 | 0.83 | 1.18 | 1.34 |
| | RBD | 0.40 | 0.34 | 0.50 | 0.64 | 0.66 | 0.96 |
| Reference (PMID) | | 32225176 | 32275855 | 32132184 | 32942285 | 33271067 | 33277323 |
